# *De novo* design of semisynthetic protein nanopores

**DOI:** 10.1101/2025.09.08.674823

**Authors:** Lee Schnaider, A. Katherine Hatstat, Alistair J. Scott, Sophia K. Tan, Richard G. Hambley, William M. Dawson, Rhys C. Griffiths, Rian C. Kormos, Arthur A. Melo, Eric Tse, Nicholas F. Polizzi, E. Jayne Wallace, Gregory E. Merz, William F. DeGrado

## Abstract

Protein nanopores are essential components of single-molecule oligonucleotide sequencing and sensing devices. Here, we demonstrate that installing additional *de novo* subunits enables large-scale architectural changes of nanopore complexes. We design *de novo* proteins that integrate seamlessly with the CsgG pore to form 18-subunit, 315-kilodalton complexes with precisely sculpted pore architectures and tailored ion conduction, opening new possibilities for engineering enhanced nanopores with customized structural and functional properties.

---

Macromolecular assemblies that form nanomachines are ubiquitous in nature and have many industrial applications^1,2^. For example, membrane-spanning nanopores are widely used in commercial nucleic acid sequencing devices, and they also show promise for protein sequencing and metabolite detection^3–5^. CsgG, a bacterial lipoprotein pore that forms a nonameric channel, has been extensively engineered for these applications^6–8^. For oligonucleotide sequencing, it was engineered to have a constriction on the order of the size of nucleotide bases to identify each base by resolving their differential effects on ion conduction as nucleotides are threaded through the channel. However, there are limitations to what can be accomplished through modification of natural proteins; for example, existing nanopore sequencing devices struggle to achieve precise sequencing in regions of repeating nucleotides^6^. *De novo* protein design provides a method to address this specific challenge and more generally enables the construction of pores of any arbitrary shape profile as required for a given application. Design of fully synthetic ion-conducting pores may eventually provide a solution to this problem^9–11^. More immediately, we suggest that *de novo* design might be used to build semisynthetic proteins that enhance existing pores engineered from natural proteins used for sequencing. The additional subunits might allow one to (i) fine-tune the overall conductivity and selectivity for permeating ions and (ii) sculpt the overall pore length and radial profile to enable reading of nucleic acids over a longer window.

Here, we sought to design a nonameric protein that docks onto CsgG-F56Q, a CsgG derivative engineered for nucleic acid sequencing^6^ (herein referred to as CsgG for simplicity), creating a cone-shaped extension of the pore that places an ∼20 Å aperture ∼50 Å from the existing CsgG constriction near the center of the membrane. Furthermore, we endeavored to design relatively small subunits (<60 residues) to allow them to be chemically synthesized, thereby facilitating easy incorporation of unnatural amino acids for future derivatization. This work focuses on the challenges associated with the design of such macromolecular complexes.

As a starting point, we used an 18-subunit pore-forming complex, CsgG:CsgF, consisting of nine CsgG subunits and nine CsgF subunits (CsgG:CsgF)_9_. In Gram-negative bacteria, CsgG forms a nonameric barrel, and a short (30-residue) segment of CsgF binds into the CsgG pore, assisting in the secretion of curli fibers^6^. CsgF binding alters conductance and base-discrimination properties, and derivatives of (CsgG:CsgF)_9_ have shown promise for dual-constriction DNA sequencing^6,12^. However, the limited stability and lack of structural programmability of the complex hindered practical applications. CsgF binds to one end of the pore as a simple extended peptide with a short helix (Fig. 1a), providing a potential anchor for a *de novo* construct that extends the pore length and modulates its geometry.

**Figure 1:**
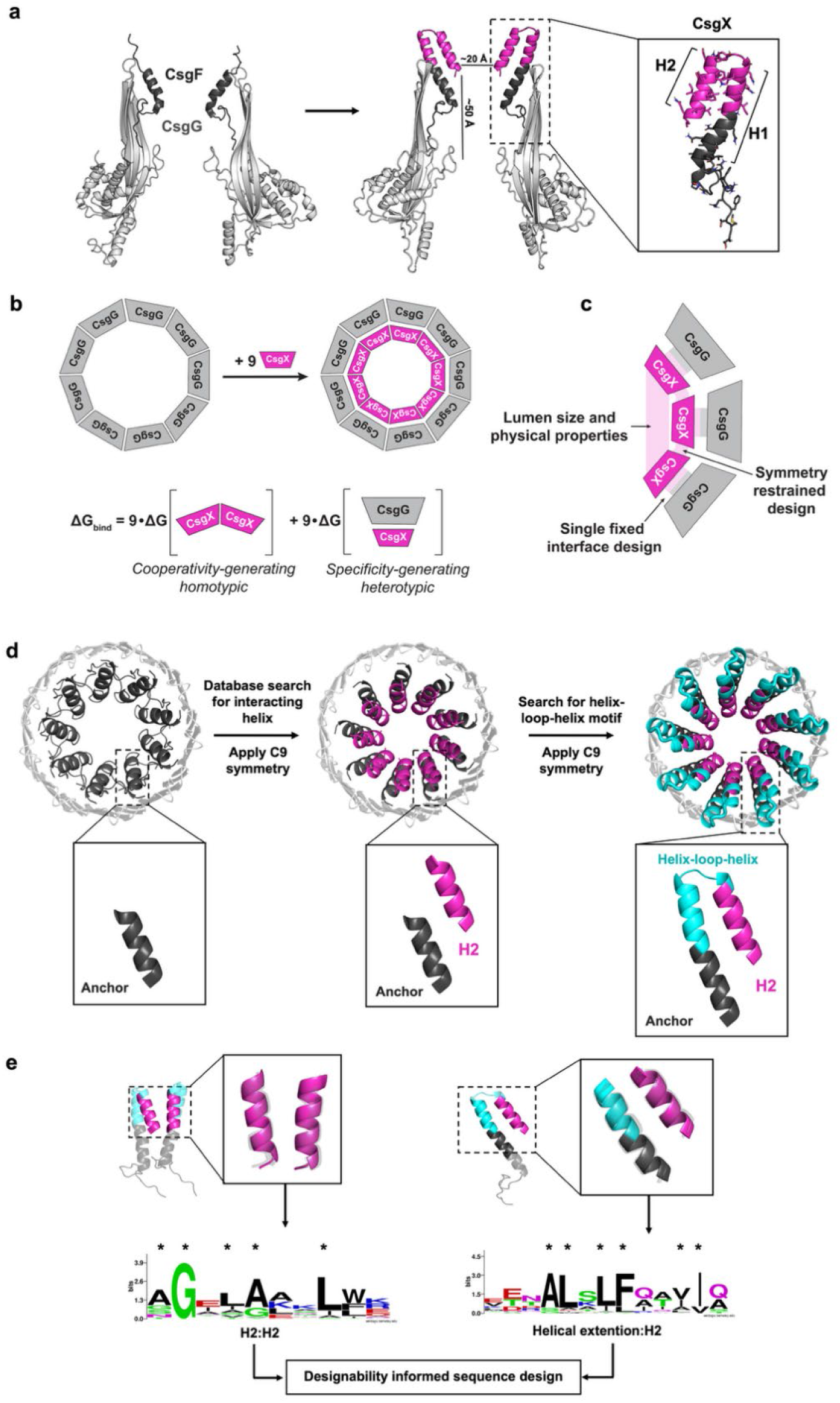
Integrated design of semisynthetic CsgG:CsgF:CsgX nanopores. **a**, Schematic of pore engineering. Starting from the (CsgG:CsgF)_9_ complex (left), a *de novo* designed helical hairpin (CsgX, magenta) is precisely grafted onto the C-terminus of CsgF (dark grey), allowing extension into the CsgG lumen (light grey). The design begins with the Anchor helix from CsgF (dark grey) and the de novo portion includes an extension of the Anchor helix to afford H1, a loop, and a second helix H2 (magenta). The boxed inset shows the resulting CsgX monomer (magenta). **b**, Schematic of pore assembly of CsgX (magenta) with CsgG_9_ (light grey). The binding energy (ΔG_bind_) is the sum of homotypic (CsgX:CsgX) and heterotypic (CsgG:CsgX) interactions over all subunits. **c**, Key structural features that needed to be accommodated during design include pore lumen dimensions, symmetry, and the fixed structure and sequence of (CsgG)_9_. **d**, Design approach. Starting from the CsgF Anchor (left), a single helix is geometrically docked against the Anchor and propagated by C9 symmetry (magenta; middle). MASTER^13^ is then used to identify geometrically compatible helical extensions of the Anchor and an adjoining helix, creating a helix–loop–helix motif (cyan; right). Candidate motifs are/were? filtered for structural clashes and a cone-like overall structure, funneling to a ∼ 20 Å restriction. **e**, The prevalence of the helical interface geometries across the PDB is evaluated using MASTER^13^ to identify frequently occurring, designable motifs. These motifs (transparent grey, overlaid on the H1:H2 and Helical extention:H2 helices) are used to generate sequence logos that capture position-specific residue preferences (right), which are considered as restraints for Rosetta-based sequence design (residue seeds are marked with an asterisk).

However, building such a protein (Fig. 1a-c) which seamlessly integrates with the existing CsgG pore is challenging as it requires (i) designing within the highly confined and structurally nonuniform pore, (ii) creating a new lumen with bespoke geometric and physicochemical properties and (iii) adhering to C9 symmetry. Moreover, to be useful for single-molecule sequencing applications, the components must assemble with complete cooperativity. Here we develop and experimentally validate a design strategy that achieves these objectives.

## Design approach

We sought to design an additional *de novo* domain that extends well beyond the (CsgG:CsgF)_9_ pore and creates a wide opening that funnels down to an 18 – 23 Å diameter (Cα to Cα) constriction 46 – 52 Å (Cα to Cα) above the main CsgG constriction, affording a highly structurally stable pore with low conductance fluctuations. To achieve these objectives, we targeted a *de novo* helix-loop-helix motif, in which the first helix (H1) packs against the CsgG barrel, while the second helix (H2) faces the pore (Fig. 1a). To ensure robust complex formation, it was essential to optimize the interactions along each interface: H1:H1, H2:H2, H1:H2 and H1:CsgG, all while maintaining C9 symmetry. This creates a challenging mutual optimization problem (Fig. 1b,c).

Backbone design was accomplished using the short 11-residue helix of WT CsgF (residues N18 – A28) as an “Anchor” from which to extend H1 and incorporate it into a hairpin (Fig. 1d, left; Supplementary Data Fig. 1). The hairpin was constructed by searching the Protein Data Bank (PDB) using the program MASTER^13^ for a helix (H2), which could interact favorably with the CsgF helix, while also forming homomeric interactions across the symmetry axis. A MASTER^13^ search identified candidates for H2, which dock against this helix in a highly prevalent geometry in the PDB (Fig. 1d, middle). We next applied C9 symmetry to generate the full assembly and eliminated structures with clashing helices and those that did not fulfill the structural requirements listed below (Fig. 1d; Supplementary Data Fig. 1). A second MASTER^13^ search was used to find a fragment that elongates the CsgF anchoring helix to complete H1, while also forming a loop that connects H1 to H2 (Fig. 1d, right; Supplementary Data Fig. 2). The application of C9 symmetry completed the design. Following each MASTER^13^ search, we filtered the output to select structures with favorable interactions across the symmetry-generated helical interfaces, as assessed by the frequency of occurrence of these geometries in the PDB (Fig. 1d,e). This metric, together with the resulting pore lumen architecture and pore diameter, serve as the criteria by which backbones were selected.

**Figure 2:**
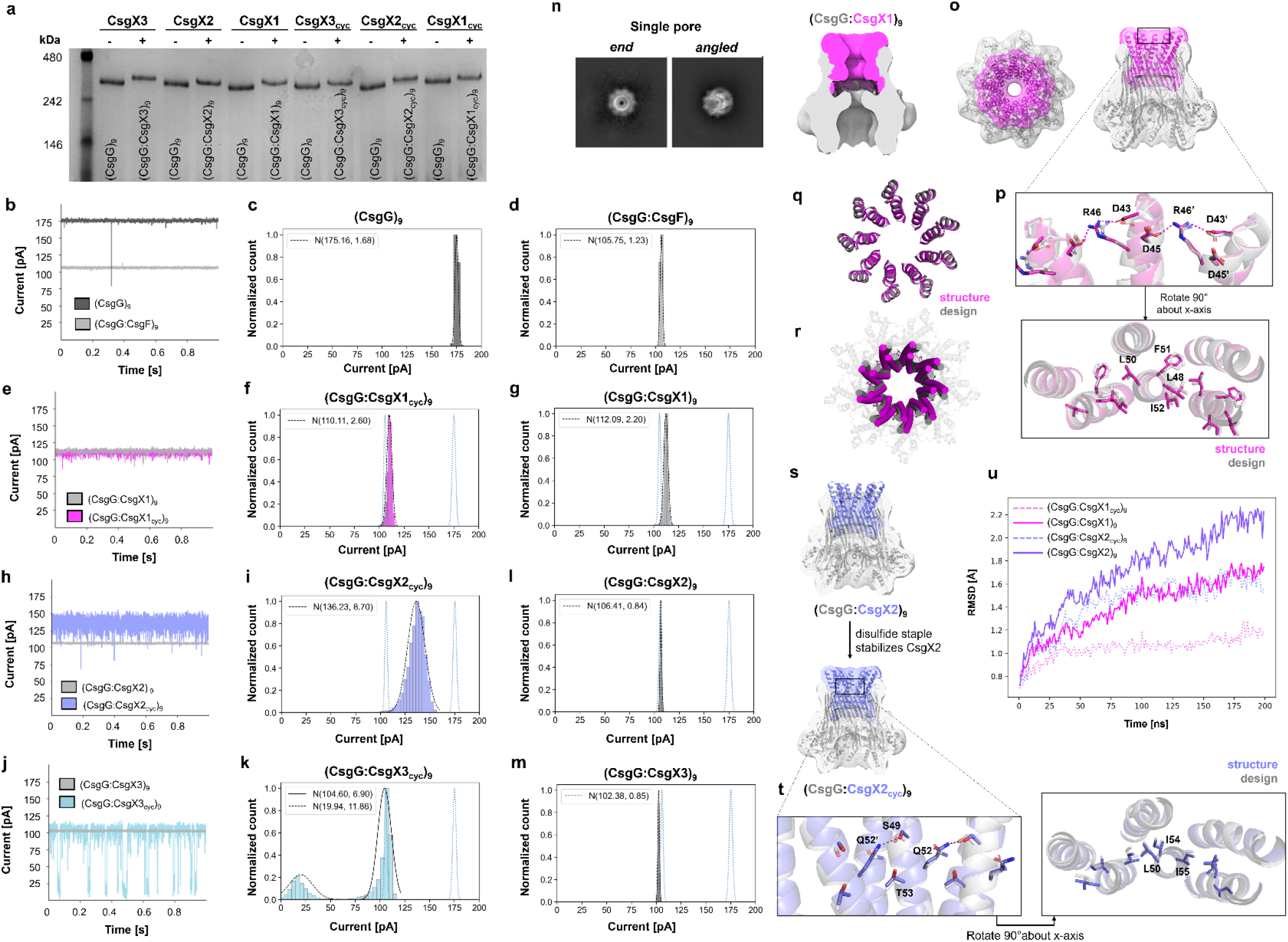
Experimental and structural characterization of CsgG:CsgX complexes. **a**, Blue native gel electrophoresis of the CsgG:CsgX assemblies confirm that all six of the CsgX variants assembled into homogeneous 9:9 complexes with CsgG, with no detectable sub-stoichiometric intermediates. **b, e, h, j**, Single-channel recordings of (CsgG:CsgX)_9_ complexes. Representative current traces at 180 mV from synthetic membrane recordings of the (CsgG:CsgX)_9_ complexes inserted into a MinION flow cell from Oxford Nanopore Technologies. Current/time traces were collected by measuring the current with a frequency of 4kHz. The individual values of the current were analyzed using a normal distribution, giving a mean current and an associated standard deviation. The theoretical curves from fitting the data for(CsgG)_9_ and (CsgG:CsgF)_9_ overlayed on each of the (CsgG:CsgX)_9_ plots in light blue dashed lines (105 pA and 175 pA, respectively) for reference (**c**,**d**, CsgG_9_ and (CsgG:CsgF)_9_, **f**,**g**, (CsgG:CsgX1_cyc_)_9_ and (CsgG:CsgX1)_9_, **i**,**l**, (CsgG:CsgX2_cyc_)_9_ and (CsgG:CsgX2)_9_, **k**,**m**, (CsgG:CsgX3_cyc_)_9_ and (CsgG:CsgX3)_9_). **n**, Representative 2D class averages (left) highlighting views of the single nanopore assembly (CsgG:CsgX1)_9_ and clipped 3D volume of the reconstructed density map for (CsgG:CsgX1)_9_, low-pass filtered to 15 Å, displayed at contour of 0.06 in UCSF ChimeraX_22_. Density for CsgG and CsgX1 are colored grey and magenta, respectively. **o**, Low pass filtered 3D volume (transparent grey/magenta) overlaid on the structural model of (CsgG:CsgX1)_9_ shows unambiguous density for the full CsgX1 hairpin and for all nine CsgX1 monomers in the assembly (end view, left; clipped side view, right). **p**, Comparison of design model (grey) to structure (magenta) of 3 CsgX1 monomers, with side chains displayed for key polar interactions (R46, D43, D45; top) and hydrophobic packing residues (L48, L50, F51, I52; bottom). **q**, Alignment of design (gray) to structure (magenta) of designed residues (31-57) of (CsgX1)_9_ show low overall RMSD (< 1 Å). **r**, In the full (CsgG:CsgX1)_9_ structure, the CsgX1 hairpin (magenta) is shifted (∼15°) about the axis of symmetry relative to the design model (gray), while retaining the overall pore geometry. **s**, 3D volumes of (CsgG:CsgX2)_9_ and (CsgG:CsgX2_cyc_)_9_, low-pass filtered to 15 Å, contoured to 0.09 in UCSF ChimeraX_22_. CsgG and CsgX2 shown in transparent grey and purple, respectively, and overlaid on (CsgG:CsgX2)_9_ design model, show that the (CsgG:CsgX2)_9_ volume has no observable density for the designed hairpin, while (CsgG:CsgX2_cyc_)_9_ has clear density for the full CsgX2. **t**, Comparison of design model (grey) to structure (purple) of 3 CsgX2_cyc_ monomers, with side chains displayed for key polar interactions (S49, Q52, T53; left) and hydrophobic packing residues (L50, I54, I55; right). **u**, RMSD of the peptide Cα atoms over a 200 ns molecular dynamics (MD) production run of the designed (CsgG:CsgX)_9_ models. The initial 20 ns of the 220 ns simulation were excluded from analysis, and RMSD values were calculated relative to the structure at 20 ns. Over the final 50 ns of the simulation, (CsgG:CsgX2)_9_ (solid purple) exhibited a higher rate of RMSD change (2.82 Å/μs) compared to the other samples (1.69 Å/μs, 2.43 Å/μs, and 1.11 Å/μs for (CsgG:CsgX1)_9_ (solid pink), (CsgG:CsgX1_cyc_)_9_ (dashed pink), and (CsgG:CsgX2_cyc_)_9_ (dashed purple), respectively).

Sequences were designed using symmetric sequence design with Rosetta^14^, which was seeded with sequence logos associated with structural features obtained from MASTER^13^ (Fig. 1e). To achieve the desired low-noise conductance, we sought to design a lumen lined with polar residues that were locked into a fixed geometry to avoid dynamic fluctuations that could give rise to noisy channel recordings. Thus, we selected sequences in which the pore-lining residues engaged in favorable electrostatic and hydrogen-bonding interactions and uniform packing of apolar residues along each interface. We chose five sequences to experimentally characterize, designated CsgX1 – CsgX5, each with a ∼27-residue *de novo* helical hairpin motif. Disulfides are often included in the design of small *de novo* proteins (<60 residues) to increase their stability and conformational rigidity^15,16^. Therefore, we also synthesized cyclic CsgX derivatives with a disulfide between residues 24 and 58, affording a total of 10 proteins (Supplementary Data Fig. 3).

## Experimental characterization

All 10 designs were found to insert into synthetic membranes and form conducting channels^6^. Only six (representing linear and cyclic versions of three sequences) gave recordings consistent with the formation of the desired (CsgG:CsgX)_9_ complex^6^, and hence these were investigated further. These six CsgX derivatives formed homogeneous 9:9 complexes with CsgG, with no evidence of sub-stoichiometric intermediates, as assessed by blue native gel electrophoresis (Fig. 2a; Supplementary Data Fig. 4). The efficiency of complex formation and lack of partially assembled complexes indicates high cooperativity of assembly, which is essential to obtain a uniform distribution of pores for electrophysiological studies.

Nanopore sequencing requires a high-conductance, 15 – 20 Å pore, with low noise occuring during collection of current/time traces. The (CsgG)_9_ pore has a uniform conduction state of 175.1 with an associated standard deviation of 1.6 pA (Fig. 2b,c) associated with the variation of the current. Incorporation of the CsgF subunits to form the (CsgG:CsgF)_9_ complex decreases the conduction to 105.2 ± 1.2 pA (Fig. 2b,d), consistent with the addition of a second 20.5 Å (Cα to Cα) radius^6^. Similarly, (CsgG:CsgX1_cyc_)_9_ and (CsgG:CsgX1)_9_ exhibit comparable conductance profiles of 110.1 ± 2.6 pA and 112.0 ± 2.2 pA, respectively, which are consistent with the designed 19.5 Å constriction, and indicate successful pore insertion and folding (Fig. 2e-g). Both (CsgG:CsgX2_cyc_)_9_ and (CsgG:CsgX3_cyc_)_9_ have conduction states distinct from (CsgG:CsgF)_9_, although the noise level is greater. Specifically, (CsgG:CsgX2cyc)_9_exhibits greater variability (136.2 ± 8.7 pA; Fig. 2h,i), while (CsgG:CsgX3cyc)_9_displays two distinct conductance states (104.6 ± 6.9 pA and 19.9 ± 11.8 pA; Fig. 2j,k). This finding suggests that they might have conformational fluctuations that occur on the micro-to milli-second time regime associated with the 4 kHz frequency of data collection. Model-building indicates that at the narrowest point of the de novo restriction, variation of the sidechain rotamers could give rise to up to 7 Å in the pore radius, possibly explaining the current fluctuation. Finally, the linear versions of these proteins, (CsgG:CsgX2)_9_ and (CsgG:CsgX3)_9_, gave traces that are indistinguishable from those of (CsgG:CsgF)_9_: 106.4 ± 0.8 pA and 102.3 ± 0.8 pA for (CsgG:CsgX2)_9_ and (CsgG:CsgX3)_9_, respectively (Fig. 2l,m). It is possible that, under these experimental conditions, the *de novo* portions of CsgX are not well folded and possibly extruded outside of the lumen.

## Structural characterization of pores

Given the favorable electrophysiological properties of (CsgG:CsgX1)_9_ and (CsgG:CsgX2)_9_, we structurally characterized the linear and cyclic forms of these assemblies using single particle cryo-electron microscopy (cryo-EM) (Fig. 2n-t). Particles from the collected micrographs for (CsgG:CsgX1)_9_ and (CsgG:CsgX2)_9_ afforded 2D class averages showing single donut-shaped particles with high-resolution features, as well as head-to-head dimers (Fig. 2n; Supplementary Data Fig. 5,6), in agreement with previously solved structures of (CsgG)_9_. ^6^From the single pore particle classes of (CsgG:CsgX1)_9_, a structure was determined to 2.6 Å resolution, which showed unambiguous density for the full-length (CsgG:CsgX1)_9_ complex (Fig. 2o; Supplementary Data Fig. 5). As in the design, the nine CsgX monomers form a funnel-like architecture, with their N-terminal portion creating a stem that inserts into the CsgG_9_ barrel, and the hairpins forming a cone-shaped extension. The (CsgG)_9_ portion of the (CsgG:CsgX1)_9_ complex is consistent with previously reported structures of CsgG in complex with CsgF (PDB: 6SI7^6^) (Cα RMSD = 2.53 Å).

The observed monomer of the *de novo* CsgX1 is in excellent agreement with the design (Cα RMSD = 0.45 Å for the *de novo* designed CsgX extension, Fig. 2q,p). The intrahelical packing between H1 and H2 in the designed portion agrees well with the design, showing the interdigitation of apolar sidechains L48, L50, F51, and I52 (Supplementary Data Fig. 7). The full network of 27 luminal polar interactions, which includes D43, D45, and R46 in each subunit, were indeed observed precisely as per the design (Fig. 2p, Supplementary Data Fig. 7).

The nonameric CsgX1 complex is inserted into CsgG_9_, but, as compared to the model, the CsgX1_9_ ring is rotated ∼15° about the axis of symmetry (Fig. 2q,r). This rotation slightly changes the docking of CsgX on CsgG (∼3 Å) but, importantly, does not affect the pore radius and lumen architecture. The structure of (CsgG:CsgX1_cyc_)_9_ is nearly identical that of (CsgX:CsgX1)_9_ (Cα RMSD of designed nonamer = 0.46 Å) (Supplementary Data Fig. 8,9), indicating that addition of the disulfide staple does not alter the behavior of the designed extension.

We also solved the structure of (CsgG:CsgX2)_9_ to 2.6 Å resolution (Supplementary Data Fig. 6). Density was observed for the entire CsgG_9_ barrel and for the first 30 residues of CsgX2_9_ (Fig. 2s, left). However, we did not observe density for residues 31-57 of CsgX2_9_. By contrast, the reconstructed map of (CsgG:CsgX2_cyc_)_9_ showed unambiguous density for the full 57 residues of CsgX2_cyc_ (Fig. 2s, lower; Supplementary Data Fig. 10). As with CsgX1 complexes, the designed homotypic and heterotypic interactions are observed experimentally with good agreement between the designed CsgX2 and the CsgX2_cyc_ model (monomer Cα RMSD = 0.65 Å over the designed hairpin; Fig. 2t). We designed a hydrogen bond network between S49, Q52, and T53 (Supplementary Data Fig. 7) but due to a rotamer shift only observed hydrogen bonds between S49 and Q52 (Fig. 2t; Supplementary Data Fig. 11). We also see good agreement for the designed hydrophobic packing that stabilizes the association of the helices with one another as well as the CsgG pore (Fig. 2t). As was seen for (CsgG:CsgX1)_9_ and (CsgG:CsgX1_cyc_)_9_, there is a slight rotation (∼15°) of the CsgX nonamers relative to the CsgG nonamer that does not affect pore lumen architecture.

Lastly, we used molecular dynamics to explore how conformational variability contributes to overall stability of the complexes. We simulated the computational designs of (CsgG:CsgX1)_9_, (CsgG:CsgX1_cyc_)_9_, (CsgG:CsgX2)_9_ and (CsgG:CsgX2_cyc_)_9_. Three independent 220 ns trajectories were run for each complex, and all four complexes maintained their overall structure throughout the full simulation (RMSD < 2 Å) (Fig. 2u). The cyclic form of both CsgX derivatives showed smaller fluctuations than their linear counterparts. The most dynamic complex, (CsgG:CsgX2)_9_, showed markedly higher RMSD throughout the simulation and had not stabilized after 200 ns. Interestingly, this correlates with our electrophysiological analysis, in which (CsgG:CsgX2)_9_ conductance was indistinguishable from that of (CsgG:CsgF)_9_, and agrees with our cryo-EM characterization in which (CsgG:CsgX2)_9_ was the only complex that did not have observable density for the full CsgX hairpin.

In this work, we address the challenge of designing predetermined pore geometry, with pinpoint accuracy, under symmetric constraints, and in a confined space. This requires the precise design of the CsgG:CsgX interface, the intermonomer interface and the size and shape of the resulting pore lumen. Thus, the entire surface of the designed protein needed to be carefully specified to enable seamless integration with the CsgG pore. Our method allowed us to isolate and separately optimize the multiple homomeric and heteromeric interfaces required for the designs. In the future, it should be possible to consider multiple structural and functional specifications in a single calculation using diffusion-based methods; although early attempts in this direction were not successful (Supplementary Data Fig. 12), these methods are continuously improving. Our findings demonstrate the potential of *de novo* design to enhance both natural and preexisting commercially valuable assemblies, creating semisynthetic nanomachines tailored for new practical devices.

## Methods

### Computational design

#### Backbone design

The backbones for the CsgX variants were generated via sequential searches of a non-redundant subset of the Protein Data Bank (PDB30, generated using MMSeqs2^17^) using MASTER^13^. The “Anchor” from which to extend H1 was defined as residues 18-28 of CsgF (PDB ID: 6L7C). The Anchor was used as the initial MASTER Backbone Search query to identify interacting helices (H2). This search consisted first of searching PDB30 for structural matches to the Anchor (a maximum RMSD of 1 Å (Cα to Cα) between the target and matched residues was allowed, see MASTER Backbone Search code in SI), and then extracting secondary structural motifs that are in the vicinity of target. The top 1,000 match ensembles were clustered by RMSD and the 100 top candidates from the largest cluster were selected, based on the number of closely related helix-helix pairs found in the database, after greedily clustering the output based on RMSD of the target region plus the discovered helix (at a threshold of 0.5 Å), affording the Anchor:H2 interface. C9 symmetry was then applied to generate the assembly, and structures with clashing helices were eliminated, resulting in 20 helical pairs, which moved forward to the designability analysis.

These helical pairs were scored for “designability” of the resulting Anchor:H2 and H2:H2 interfaces via a MASTER search. Clustering of the fragments was based on the combined RMSD of each of the helical pairs (RMSD < 1.5 Å, see MASTER Designability Analysis code in SI). Any symmetric assemblies with steric clashes (van der Waals overlaps of heavy atom distance < 3.5 Å) or undesirable pore lumen geometry (lumen diameter < 19 Å or > 23Å, as measured by Cα to Cα distances of the lumen-facing residues at the most occluded cross-section) were discarded.

The resulting Anchor:H2 interface (3 helical pairs in total) was used as the query for a subsequent MASTER search to extend the Anchor and install an interhelical loop, affording the CsgX hairpin (see MASTER loop building code in SI). Loop insertion lengths of 8-21 residues were evaluated. Good geometric fits were observed only with an insertion length (wgap) of either 8 or 14 residues. Candidate monomers were propagated around the C9 symmetry axis, affording 6 nonameric assemblies for assessment. These were scored for their “designability”, in the same manner as described in the above (see MASTER Designability Analysis in the SI). After filtering, the top scoring backbone was advanced to sequence design.

#### Sequence design

Sequence design was conducted with Rosetta symmetric sequence design (see Rosetta sequence design file in SI) which was seeded with residue-level constraints derived from sequence logos associated with structural features obtained from the MASTER match alignments (for residues in positions 31, 32, 34, 35, 38, 47, 48, 50, 51, 54 and 55, see Rosetta resfile in SI).

1,000 unique sequence designs were initially filtered down to 100 candidates based on lowest Rosetta energy score (< -8,200) and highest PackStat score (> 0.55). This filtered subset was further parsed to five final candidates based on (i) the presence of hydrogen bonding and/or salt bridge interactions among lumen facing side chains (as designed for the residues in positions 43, 45, and 46 in CsgX1 and 49, 52, and 53 in CsgX2 and CsgX3) and (ii) apolar packing at the helical interfaces. For these final five design candidates, a “cyclic” derivative was generated by mutating residue 24 to cysteine, extending the C-terminal of CsgX, and installing a new C-terminal cysteine at position 58. Positions 24 and 58 were chosen because introducing cysteines at these sites allows for disulfide bond formation with favorable geometry and dihedral angles. A total of 10 CsgX variants (5 linear and 5 corresponding cyclic derivatives) were thus choosen for experimental testing (Supplemental Table 1).

### *E. coli* CsgG Pore Production

Recombinant expression vectors encoding the CsgG-F56Q variant nanopores with a C-terminal StrepII affinity tag and ampicillin resistance gene were transformed into chemically competent *E. coli* cells. The cells were plated onto an LB agar plate containing appropriate antibiotics for selection and incubated overnight at 37^°^C. LB Media with appropriate antibiotics was inoculated with a single colony from the agar plate and grown overnight at 37°C with shaking. The culture was diluted into autoinduction media plus necessary antibiotics and incubated at 18°C for 68 hours with shaking. The cells were harvested through centrifugation before being lysed and extracted into a buffer containing 1x Bugbuster extraction reagent (Merck 70921) and 0.1% DDM. The lysate was spun down (39,000 rcf) and the pore was purified using affinity chromatography (Cytiva StrepTrap™ XT, 5 mL column), heat treatment and then size exclusion chromatography (Cytiva Superdex® 200 Increase 10/300 GL), selecting for oligomeric nanopores, as determined by SDS-PAGE.

### CsgG-CsgX Complex Formation Protocol

(CsgG:CsgX)_9_ complexes were prepared from nanopores purified as above. Peptides were synthesized by Fmoc solid-phase peptide synthesis on a microwave-assisted CEM Liberty Blue™ peptide synthesiser. General synthetic procedures for Fmoc deprotection and coupling were carried out in accordance with the manufacturer’s specifications. Peptides of cyclic CsgX variants were subject to treatment with TMAD (10 eq. in pure DMSO) to induce cyclisation via disulfide bond formation, and all peptides were confirmed as the target mass using an Agilent 1260 LCMS single quad system. Nanopores were incubated with an 8x molar excess of peptide to CsgG monomer for 1 hour at 25°C. The sample was then heated at 60°C for 15 minutes.

### Blue Native PAGE

20 µL of 0.1 mg/ml (CsgG:CsgX)_9_ complex were mixed 1:1 with Native Sample Buffer (Bio-Rad). These were loaded onto a gel (Bio-Rad 4–20% Criterion™ TGX™), with Tris-Glycine (TG) buffer in the anode compartment and TG with NativePAGE™ Cathode Buffer Additive (Bio-Rad) in the cathode compartment. A protein standard (NativeMark™, Invitrogen) was included in the leftmost lane. The gel was run at 100 V for 30 minutes, after which the cathode buffer was replaced with TG. The gel was then run for a further 60 minutes at 180 mV. The gel was stained with Coomassie G-250 prior to imaging.

### Electrical Measurements

Electrical measurements were acquired for CsgG_9_ and (CsgG:CsgX)_9_ complexes that were inserted into MinION flow cells. After a single pore inserted from the *cis* side of the membrane, the *cis* compartment was perfused with 1 mL of of storage buffer (S), to remove any excess nanopores. Electrical measurements were acquired using MinION Mk1b from Oxford Nanopore Technologies. For IV curves, voltages were incrementally increased by 25 mV in alternating positive and negative potentials from 0 mV to +/-200 mV. Additional open-pore current measurements were performed at 180 mV by replacing *cis* compartment storage buffer with LSK114 sequencing buffer mixture. Here, the flowcell was first flushed with flowcell flush mixture (975 μl FCF, 25 μl FCT), then sequencing mixture (37.5 μl SB, 25.5 μl LIB, 12 μl EB) was added *via* the Spot-ON port.

Note that in all recordings, a positive current is one in which cations move from *trans* to *cis*. Raw data were collected in a bulk FAST5 file using MinKNOW software (Oxford Nanopore Technologies) and exported to CSV format for plotting.

### Electrophysiology data analysis

Current/time traces from single-channel recordings were extracted and baseline-corrected for comparative visualization. For each design, histograms of open-pore current values were constructed using 30 equally spaced bins spanning the observed current range. Histogram counts were normalized by the height of the tallest bin, such that each histogram had a maximum bar height of 1.

To estimate the characteristic open-pore conductance of each design, Gaussian functions were fit to the unbinned current values using maximum likelihood estimation of the mean (μ) and standard deviation (σ). The resulting probability density functions were evaluated over 500 linearly spaced points and normalized to unit peak height to match the histogram scaling. In the case of (CsgG:CsgX3_cyc_)_9_, a two-component Gaussian mixture model was fit, and the component curves were rescaled.All analyses were performed using Python 3.11 with the numpy, matplotlib, scipy, and scikit-learn libraries.

### Cryo-EM Sample Preparation and Data Collection

(CsgG:CsgX1)_9_, (CsgG:CsgX1_cyc_)_9_, (CsgG:CsgX2)_9_ and (CsgG:CsgX2_cyc_)_9_ (3 μL) at concentration of 0.3 mg/mL was added to 200 mesh 1.2/1.3 Au Quantifoil grids coated with a layer of graphene oxide^18^ which had not been glow discharged. After 15 seconds, grids were blotted for 7 s at 4°C and 100% humidity using an FEI Vitrobot Mark IV, followed by plunge freezing in liquid ethane. Super-resolution movies were collected at a nominal magnification of 105,000x (physical pixel size: 0.417 Å/pixel) on a Titan Krios (Thermo Fisher Scientific) operated at 300 kV and equipped with a K3 direct electron detector and BioQuantum energy filter (Gatan, Inc.) set to a slit width of 20 eV. A defocus range of -0.8 to -1.8 μm was used, with a total exposure time of 2.024 seconds fractionated into 80 0.025-second subframes. The total dose for each movie was 46 electrons/Å^2^. Movies were motion corrected using MotionCor2^19^ in Scipion^20^ and were Fourier cropped by a factor of 2 to a final pixel size of 0.834 Å/pixel.

For each dataset, all processing was done in CryoSPARC v4.5^21^. For particle picking, blob picker was set to 110 Å minimum and 220 Å maximum diameter. Particles were inspected before extracting with a box size of 440 unbinned pixels. Multiple rounds of 2D classification were performed, and good classes were selected for continued processing. Two volumes were generated *ab initio* and used as starting models for 3D classification. The larger class with clear features consistent with a nanopore structure was selected and homogeneous and non-uniform refinements were run on these particles (see Table S2).

For (CsgG:CsgX1_cyc_)_9_, the final particle stack was symmetry expanded in CryoSPARC using C9 symmetry. Local refinement, using a mask encompassing the entire complex, was performed on this particle stack to give a final map with 2.7 Å resolution.

### Molecular Modeling

The design model of (CsgG:CsgX1)_9_ nonamer was used as an initial model and docked into the EM map using UCSF ChimeraX’s fit in map function^22^. Refinement was done in Phenix using RealSpace Refine^23^, followed by local refinement in Isolde^24^.

### Molecular Dynamics Simulations

Three independent molecular dynamics simulations were performed, starting with the computational design. The linear and cyclic versions of (CsgG:CsgX1)_9_ and (CsgG:CsgX2)_9_ were modeled with C-terminal amidation to reflect their experimentally synthesized forms. All simulations were performed using the AMBER22^25^ with the ff19SB force field^26^ and the OPC water model^27,28^. To build the simulation box, each structure was first solvated in a rectangular periodic box with 8 Å padding from the nanopore complex, and ionized with Na+ and Cl-ions to match buffer concentrations used in experimental assays. For simplicity, the nanopore was solvated in water, with the transmembrane atoms restrained using a harmonic potential with a 15 kcal/(mol·Å^2^) force constant to mimic membrane anchoring. Hydrogen-containing bonds were restrained using the SHAKE algorithm^29,30^. Long-range electrostatics were computed using the particle mesh Ewald method^31^, and nonbonded interactions were truncated at 12 Å.

Three independent 220 ns trajectories were run for the simulation boxes (each comprising 84,980-86,308 heavy atoms), and the first 20 ns were excluded; only the final 200 ns (i.e., the “production run”) were analyzed. Simulations began with 1,000 restrained steepest descent minimization steps, followed by a maximum of 7,000 steps using the conjugate gradient method. The systems were then heated from 100 to 293 K over 50 ps in the NVT ensemble with a Langevin thermostat and a 1 fs time step, followed by pressure equilibration at 1 atm using the Monte Carlo barostat^32^ in the NPT ensemble. Throughout these minimization and equilibration steps, the nanopore complex was restrained with harmonic potentials at 15 kcal/(mol·Å^2^) initially and ramped down to 0 kcal/(mol·Å^2^) over 7 equilibration steps totaling 700 ps, with the exception of the transmembrane region, which remained restrained throughout the entire simulation. Following minimization and equilibration, each trajectory ran for 220 ns under periodic boundary conditions with 2 fs time steps. The first 20 ns of each trajectory were disregarded, and RMSDs were calculated for peptide Cα atoms relative to their coordinates at the 20 ns time point (the “production run”) and averaged across the triplicate trajectories.

## Data Availability

Scripts for protein design are provided in the Supplementary Information. The structural models are accessible via the Protein Data Bank with the following IDs: (CsgG:CsgX1)_9_, 9PNA; (CsgG:CsgX1_cyc_)_9_, 9PN8; (CsgG:CsgX2)_9_, 9PN9; (CsgG:CsgX2_cyc_)_9_, 9PNB.

## Ethics declarations

The authors declare the following competing financial interest: L.S, A.J.S, R.G.H, R.C.G, N.F.P, E.J.W, and W.F.D are co-inventors of the international patent application WO 2024/033447A1 entitled: De novo pores. A.J.S, R.G.H, R.C.G and E.J.W are members of Oxford Nanopore Technologies plc, which commercializes nanopore devices. W.F.D received financial support from Oxford Nanopore Technologies plc for this work.

## Acknowledgments

We thank Nam Hyeong Kim, Henry R. Scott and Kehan Chen for their helpful discussions and input on the manuscript. This work was supported by NIGMS F32GM143869-01A1 (to L.S.), NIGMS F32GM147962 (to A.K.H) and N IH R35 grant GM122603 (to W.F.D). This work was partially supported by NSF grant 2108660 (to W.F.D).

