## Supplemental Information for "*De novo* design of semisynthetic protein nanopores"

[1]Department of Pharmaceutical Chemistry, University of California, San Francisco, San Francisco, USA, [2]Cardiovascular Research Institute, University of California, San Francisco, San Francisco, USA, [3]Oxford Nanopore Technologies plc, Oxford, UK, [4]Institute for Neurodegenerative Diseases, University of California San Francisco, San Francisco, USA [5]Department of Neurology, University of California San Francisco, San Francisco, USA. [6]Department of Cancer Biology, Dana-Farber Cancer Institute, Boston, USA. [7]Department of Biological Chemistry and Molecular Pharmacology, Harvard Medical School, Boston, USA.

**Supplementary Data Figures** **Page**

1. Backbone generation of H2 1
2. Backbone generation of H1 extension and interhelical loop 3
3. Models of linear and cyclized CsgX variant 4
4. Full Blue native PAGE gel 5
5. Overview of structural validation of $(CsgG{:}CsgX1)_9$ 6
6. Overview of structural validation of $(CsgG{:}CsgX2)_9$ 7
7. Comparison of the inter- and intra-helical hydrogen bonding networks designed into $(CsgG{:}CsgX1)_9$ and $(CsgG{:}CsgX2)_9$ 8
8. Overview of structural validation of $(CsgG{:}CsgX1_{cyc})_9$ 9
9. Comparison of $(CsgG{:}CsgX1)_9$ and $(CsgG{:}CsgX1_{cyc})_9$ 10
10. Overview of structural validation of $(CsgG{:}CsgX2_{cyc})_9$ 11
11. Key polar interactions are shifted in the structure of $(CsgG{:}CsgX2_{cyc})_9$ relative to the design model 12
12. Design attempts using RFdiffusion to install CsgX extension 13

**Supplementary Data Tables**

1. CsgG, CsgF and CsgX sequences 15
2. Cryo-EM validation data 16

**Supplementary Scripts** 17

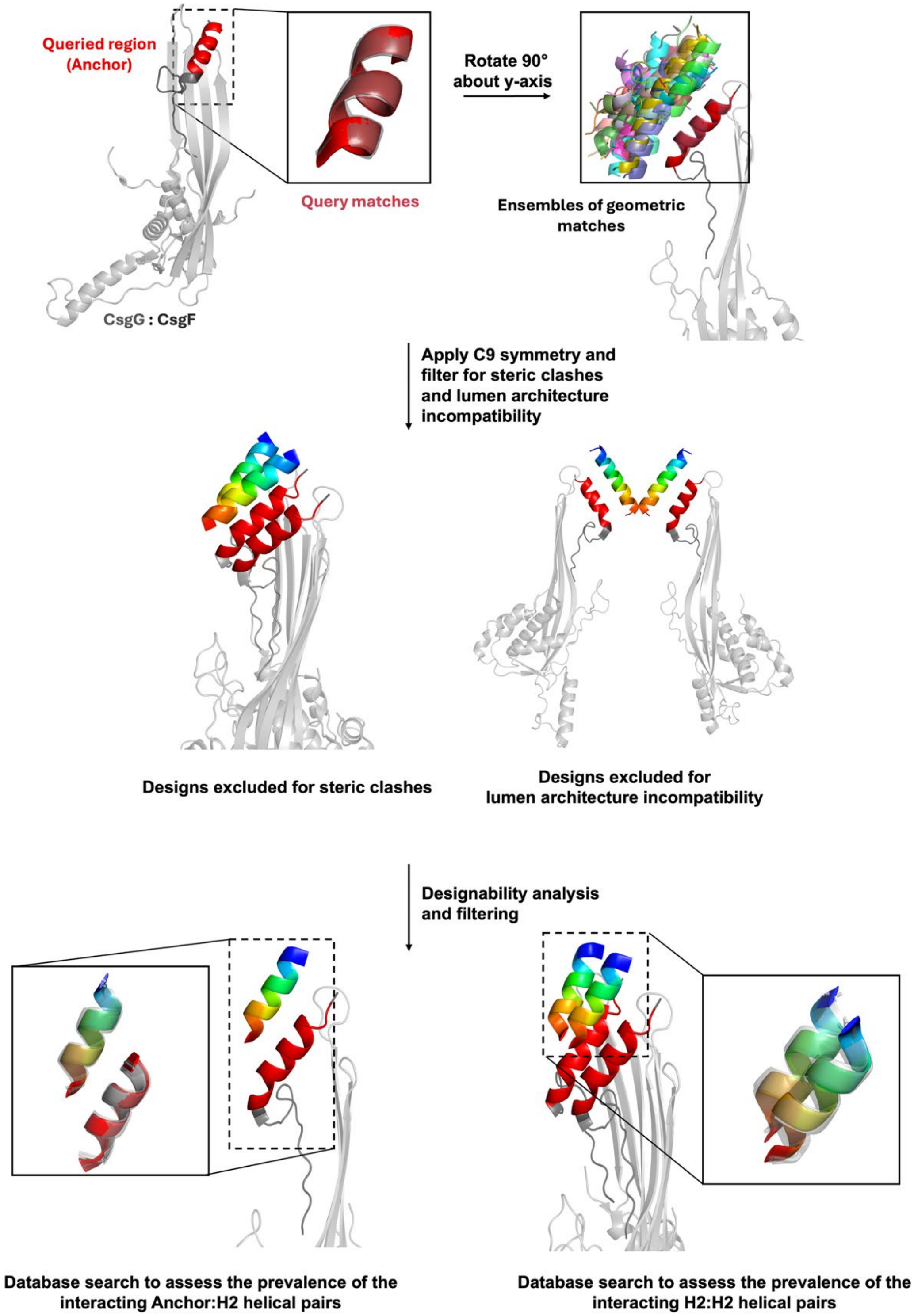


**Supplementary Data Figure 1: Design approach for H2 backbone generation using MASTER.** Backbones were generated by sequential MASTER searches of a non-redundant PDB30 dataset. The anchor helix (residues 18–28 of CsgF, shown in red; PDB ID: 6L7C) was used to identify nearby interacting helices (H2), affording 1,000 structural matches. These were clustered by RMSD, and 100 Anchor:H2 pairs from the largest cluster were selected based on geometric recurrence. C9 symmetry was applied to generate full assemblies, and 20 clash-free models with appropriate lumen geometry (19 – 23 Å diameter) were advanced to designability scoring. Representative H2 hit after symmetry was applied is shown in rainbow. Anchor:H2 and H2:H2 interfaces were evaluated via MASTER designability analysis, and three backbone models were chosen for loop building based on structural prevalence, interface geometry and structural feasibility (models with poor packing or steric occlusion were excluded).

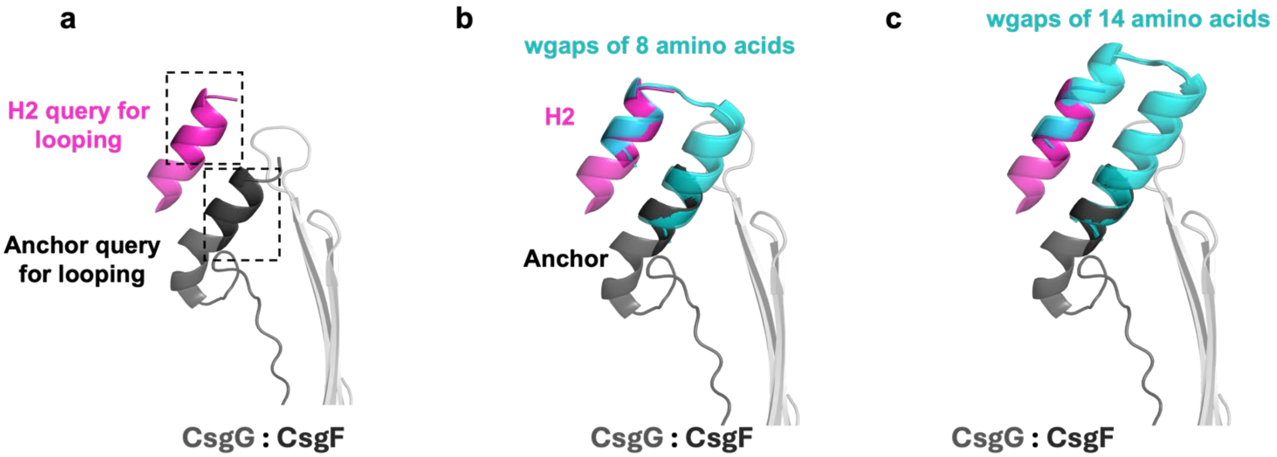


**Supplementary Data Figure 2: Design of the helix-loop-helix motif using MASTER.** Following identification of the Anchor:H2 interface, the two helices were connected using MASTER. **a,** The queried regions of the Anchor (dark grey) and H2 (magenta) are highlighted. **b,c,** wgaps (the number of residues allowed between the Anchor and H2, cyan) of 8-21 residues were evaluated, and the wgaps (the member of the largest cluster with the lowest RMSD to the queried region) with lengths of 8 amino acids (**b**) and 14 amino acids (**c**) were chosen for further designability analysis.

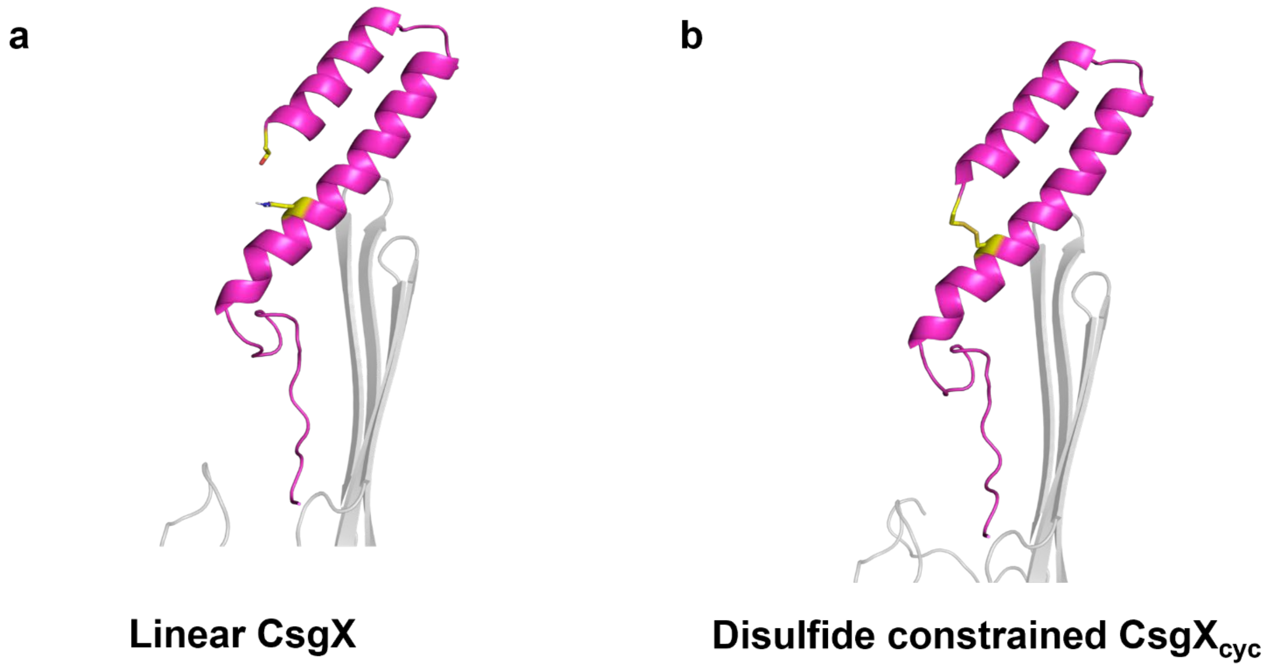


**Supplementary Data Figure 3: Design of linear (a) and cyclic (b) versions of CsgX.** Disulfide constrained CsgX peptides were designed by adding an additional C-terminal Cys residue at position 58, and mutating Asn24 to Cys.

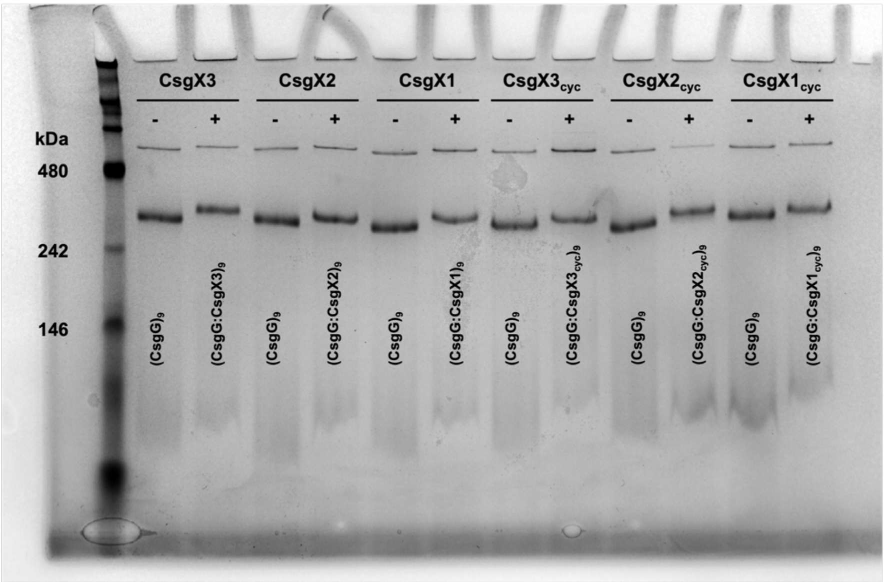


**Supplementary Data Figure 4:** Full Blue native PAGE gel for $(CsgG:CsgX)_9$ complex formation analysis. Gel image is provided as the full, uncropped image of the gel presented in Figure 2a.

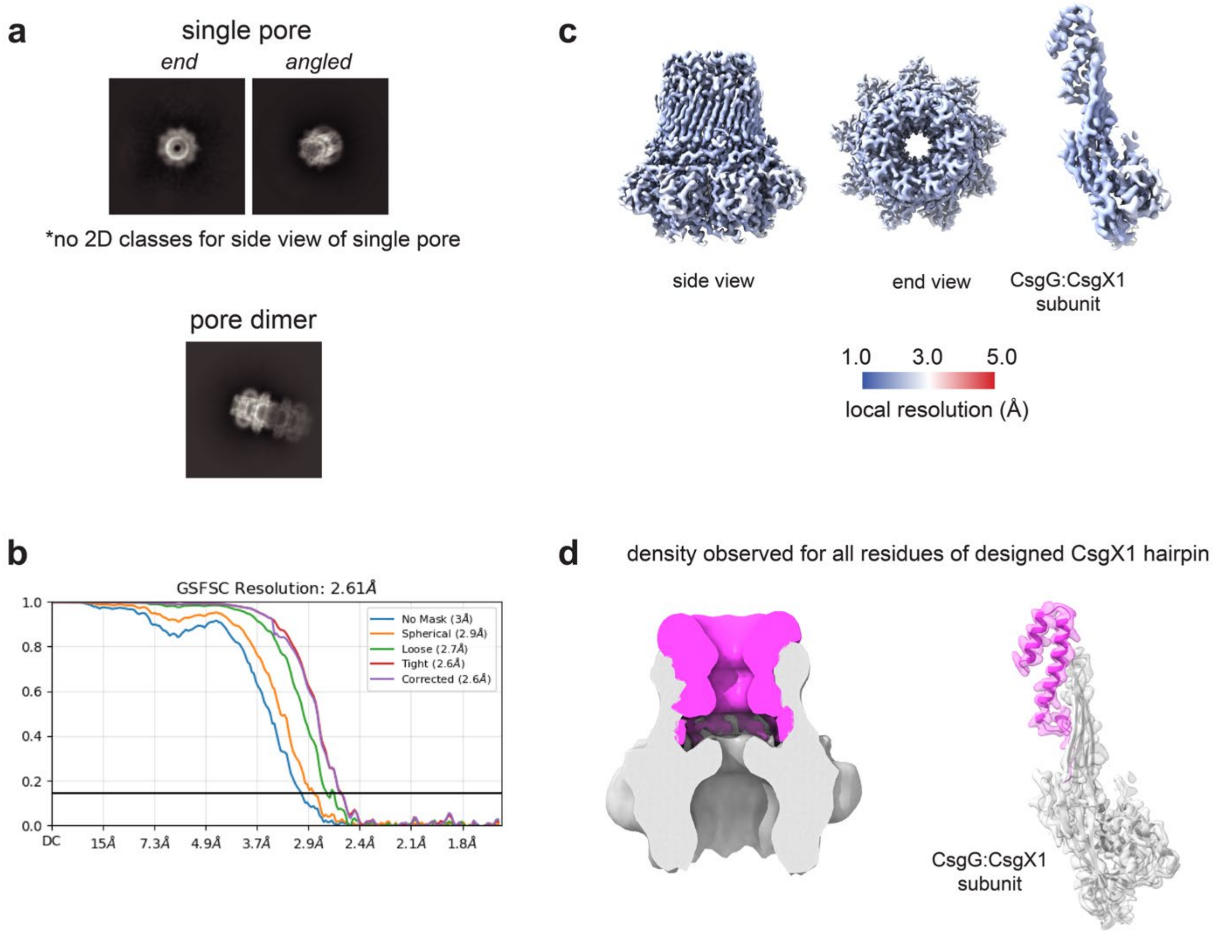


**Supplementary Data Figure 5: Overview of structural validation of (CsgG:CsgX1)$_9$. a,** Representative 2D class averages of (CsgG:CsgX1)$_9$ show a mixture of single pores and pore dimers, with varied views. **b,** Gold-standard Fourier Shell Correlation (GSFSC) curves for two independently refined cryo-EM half maps for (CsgG:CsgX1)$_9$. **c,** Local resolution of (CsgG:CsgX1)$_9$ assembly (side view, left; end view, center) and CsgG:CsgX1 subunit (right, enlarged), colored from 1.0 Å (blue) to 5.0 Å (red) resolution. **d,** Low-pass filtered (15 Å) density for (CsgG:CsgX1)$_9$ assembly (left) and density with structural model for single CsgX:CsgX1 subunit show observable density observed for all residues of CsgX1. Single pore 2D class averages and low-pass filter map are duplicated from Figure 2n.

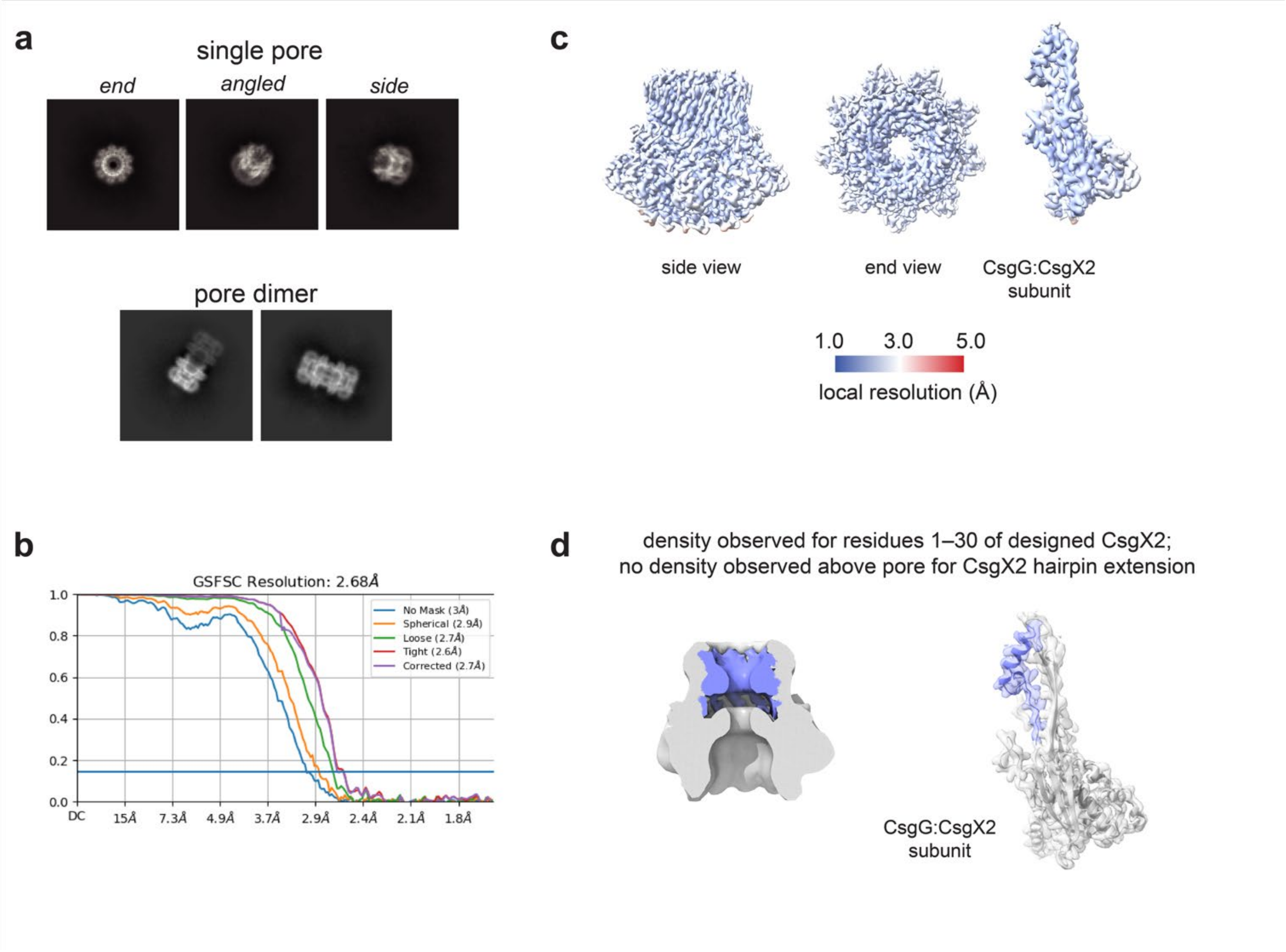


**Supplementary Data Figure 6: Overview of structural validation of (CsgG:CsgX2)$_9$ a,** Representative 2D class averages of (CsgG:CsgX2)$_9$ show a mixture of single pores and pore dimers, with varied views. **b,** Gold-standard Fourier Shell Correlation (GSFSC) curves for two independently refined cryo-EM half maps for (CsgG:CsgX2)$_9$. **c,** Local resolution of (CsgG:CsgX2)$_9$ assembly (side view, left; end view, center) and CsgG:CsgX2 subunit (right, enlarged), colored from 1.0 Å (blue) to 5 Å (red) resolution. **d,** Low-pass filtered (15 Å) density for (CsgG:CsgX2)$_9$ assembly (left) and density with structural model for single CsgX:CsgX2 subunit show no observable density observed beyond residue 30 of CsgX2, precluding structural elucidation of the designed helix-turn-helix hairpin.

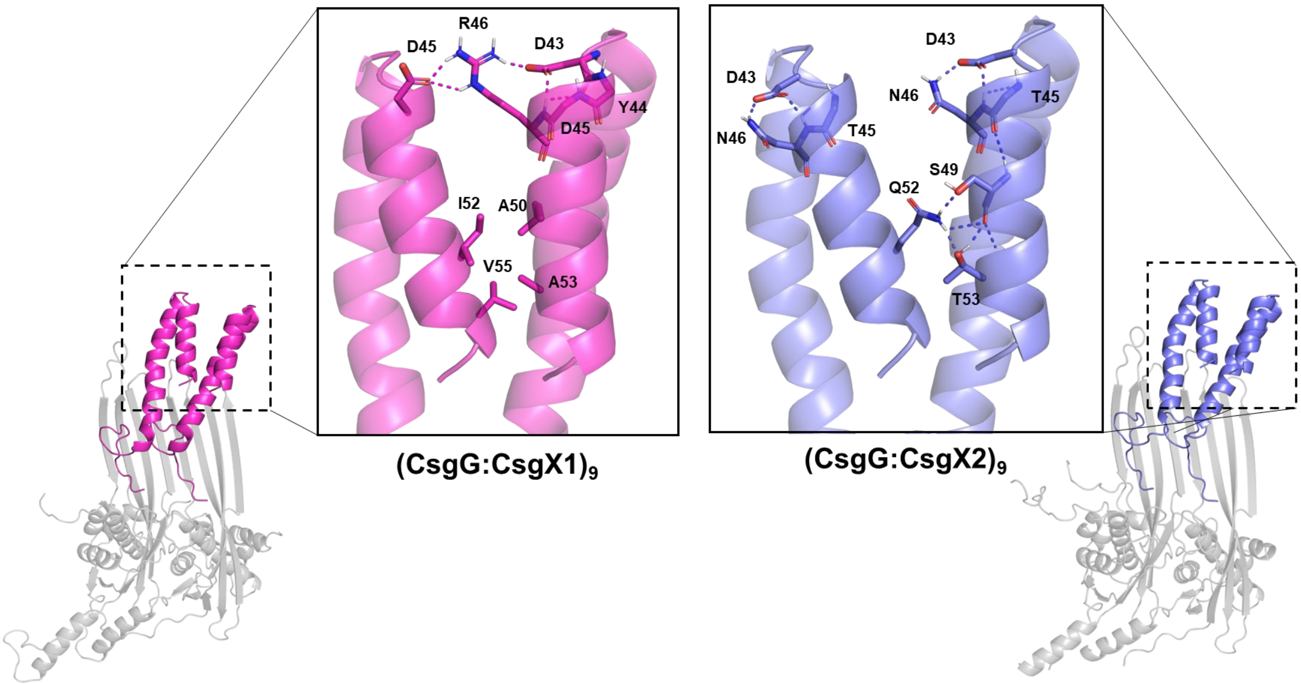


**Supplementary Data Figure 7: Comparison of the inter and intra-helical hydrogen bonding networks in design models of (CsgG:CsgX1)$_9$ and (CsgG:CsgX2)$_9$.** The computational design of (CsgG:CsgX1)$_9$ is shown in magenta (left) and the computational design of (CsgG:CsgX2)$_9$ is shown in purple (right). In (CsgG:CsgX1)$_9$ intermolecular hydrogen bonding networks were designed into the helix-loop-helix motif and hydrophobic interactions were designed into the C-terminal region of CsgX1 (left). In contrast, intramolecular hydrogen bonding networks were designed into the helix-loop-helix motif of CsgX2, and intermolecular hydrogen bonding networks were designed into the C-terminal region of CsgX2 (right).

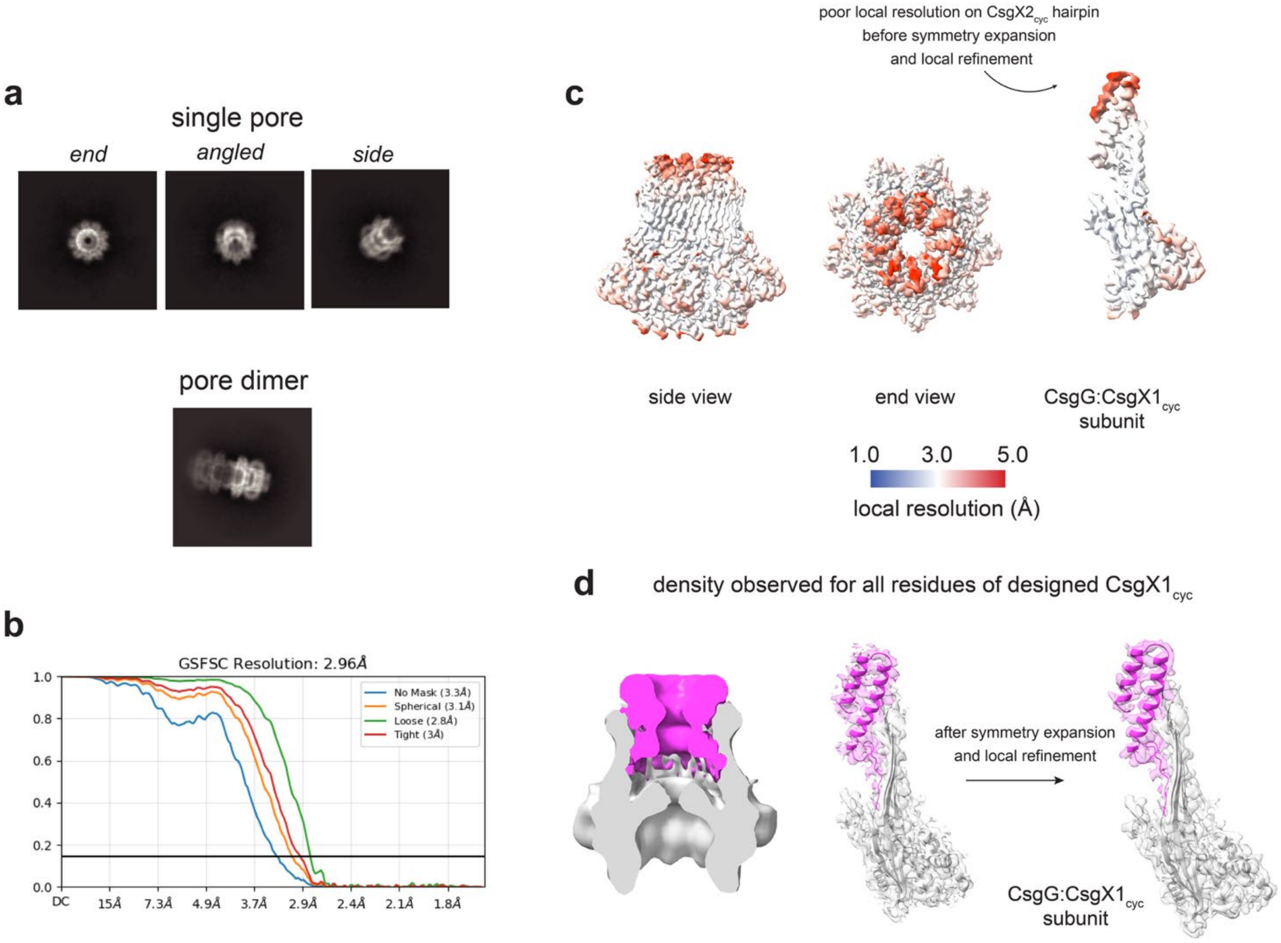


**Supplementary Data Figure 8: Overview of structural validation of (CsgG:CsgX1$_{cyc}$)$_9$. a,** Representative 2D class averages of (CsgG:CsgX1$_{cyc}$)$_9$ show a mixture of single pores and pore dimers, with varied views. **b,** Gold-standard Fourier Shell Correlation (GSFSC) curves for two independently refined cryo-EM half maps for (CsgG:CsgX1$_{cyc}$)$_9$ after symmetry expansion and local refinement. **c,** Local resolution of (CsgG:CsgX1$_{cyc}$)$_9$ assembly (side view, left; end view, center) and CsgG:CsgX1 subunit (right, enlarged). Map (before symmetry expansion and local refinement) is contoured to 0.3 and colored from 1.0 Å (blue) to 5 Å (red) resolution. **d,** Low-pass filtered (15 Å) density for (CsgG:CsgX1$_{cyc}$)$_9$ assembly (left) and density with structural model for single CsgX:CsgX1$_{cyc}$ subunit show density for full CsgX1$_{cyc}$ hairpin, with improved density after symmetry expansion and local refinement (both maps zoned around single CsgG:CsgX1$_{cyc}$ subunit at 1.80 Å radius and displayed at contour level of 0.25).

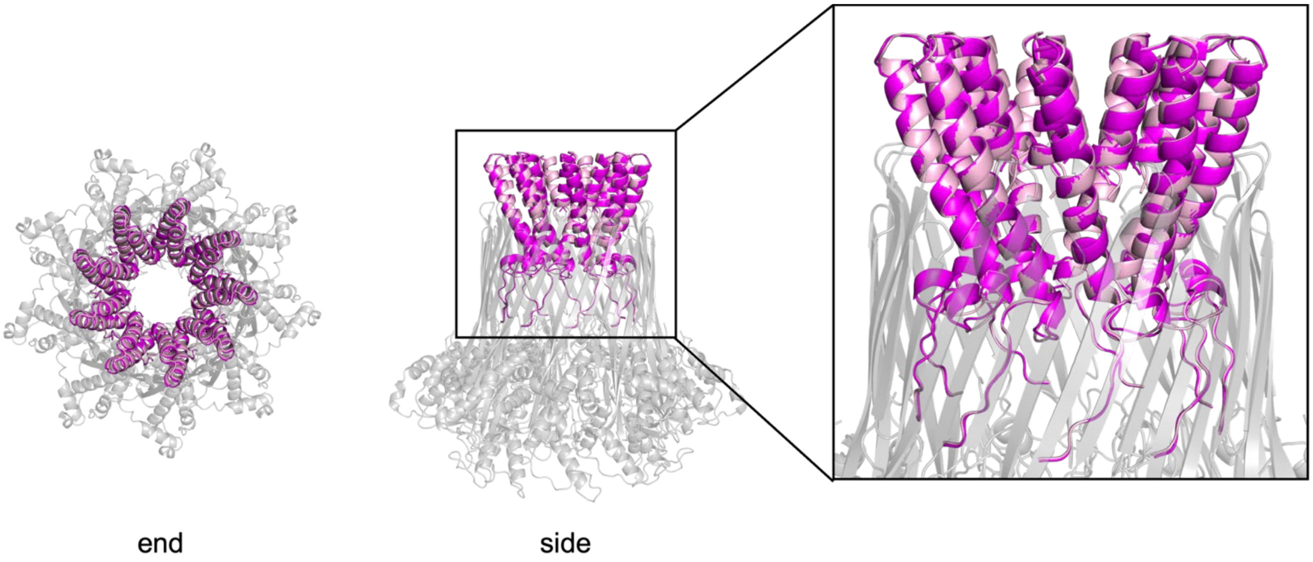


**Supplementary Data Figure 9: Addition of disulfide staple does not disrupt the overall architecture of the (CsgG:CsgX1)$_9$.** (CsgX1)$_9$ (magenta) and (CsgX1$_{cyc}$)$_9$ (light pink) have same overall topology and orientation in the (CsgG:CsgX1)$_9$ and (CsgG:CsgX1$_{cyc}$)$_9$ assemblies, respectively.

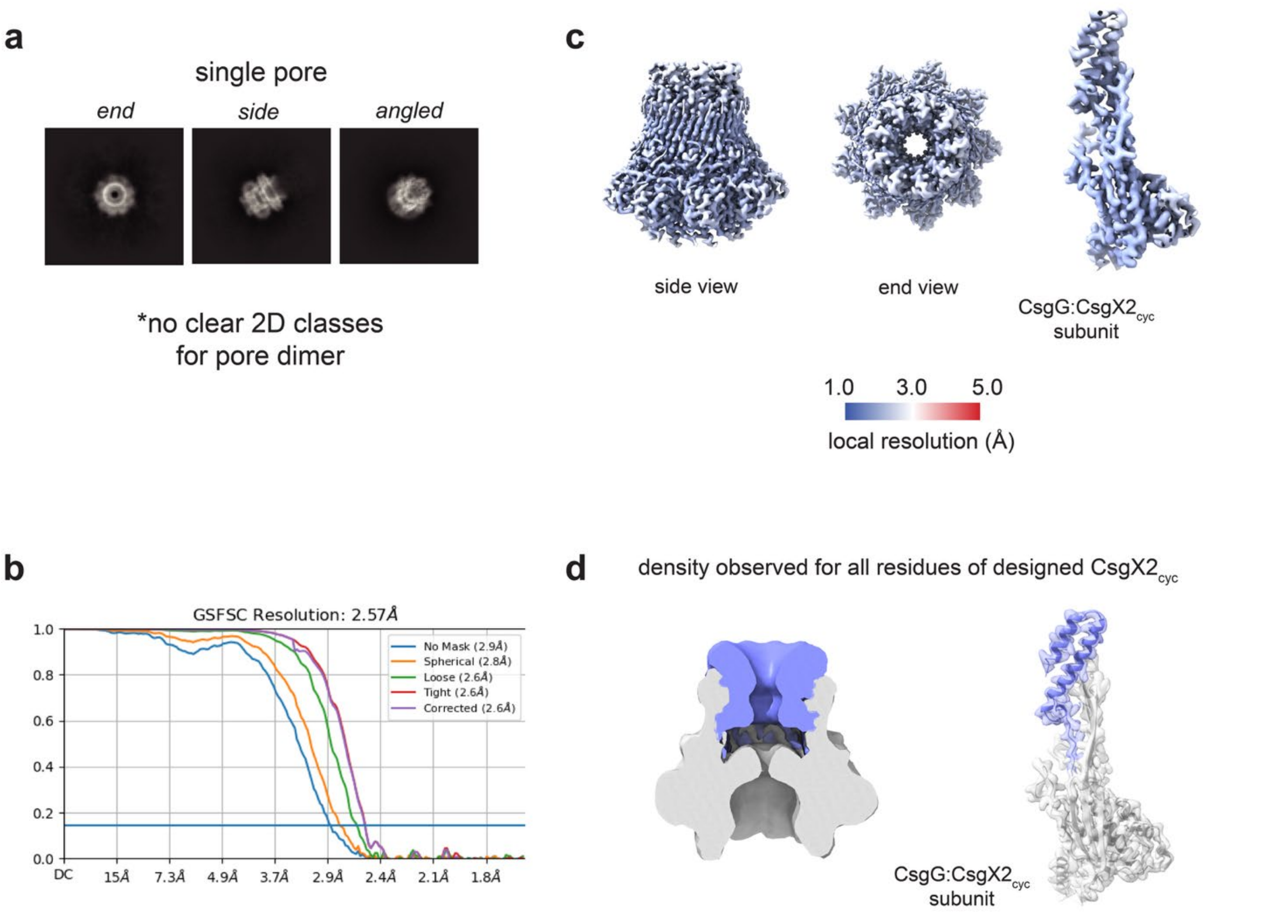


**Supplementary Data Figure 10: Overview of structural validation of (CsgG:CsgX2$_{cyc}$)$_9$. a,** Representative 2D class averages of (CsgG:CsgX2$_{cyc}$)$_9$ show single pores with varied views, but there were no 2D class averages for dimer of pores as seen with other (CsgG:CsgX2)$_9$ complexes. **b,** Gold-standard Fourier Shell Correlation (GSFSC) curves two independently refined cryo-EM half maps for (CsgG:CsgX2$_{cyc}$)$_9$. **c,** Local resolution of (CsgG:CsgX2$_{cyc}$)$_9$ assembly (side view, left; end view, center) and CsgG:CsgX2C subunit (right, enlarged), colored from 2.5 Å (blue) to 5 Å (red) resolution. **d,** Low-pass filtered (15 Å) density for (CsgG:CsgX2$_{cyc}$)$_9$ assembly (left) and density with structural model for single CsgX:CsgX2$_{cyc}$ subunit show observable density for the full designed CsgX2$_{cyc}$ hairpin.

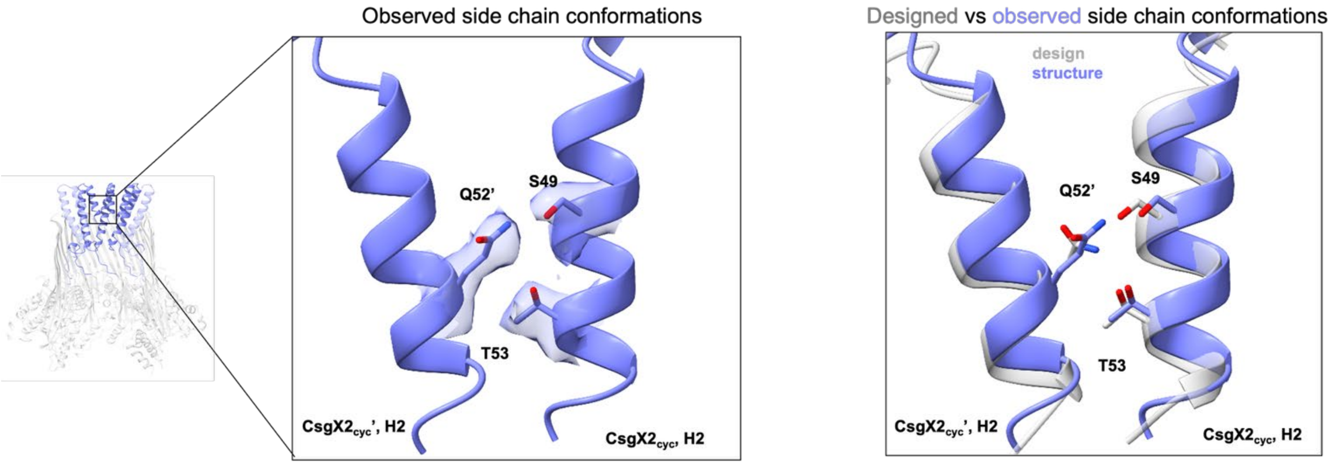


**Supplementary Data Figure 11**: **Key polar interactions are shifted in the structure of (CsgG:CsgX2$_{cyc}$)$_9$ relative to the design model**. The structure of two CsgX2$_{cyc}$ subunits, CsgX2$_{cyc}$ and CsgX2$_{cyc}$' (purple) from the full (CsgG:CsgX2$_{cyc}$)$_9$ structure are shown fit in the sharpened, refined 3D volume for (CsgG:CsgX2C)$_9$ (zoned around CsgX2C at 1.9 radius around key residues and contoured to 0.5) (left). The corresponding CsgX2 subunits from the design model (grey) are aligned with the structure displayed in purple (right). Side chains of key polar residues (S49, Q52, T53) in the refined structure fit in the observed density (left) but deviate from the designed rotamers for S49 and Q52 (right). In the experimental structure (purple) the shift in the rotamers of S49 and Q52 maintains a hydrogen bond between S49 and Q52 but disrupts the designed hydrogen bond between Q52 and T53.

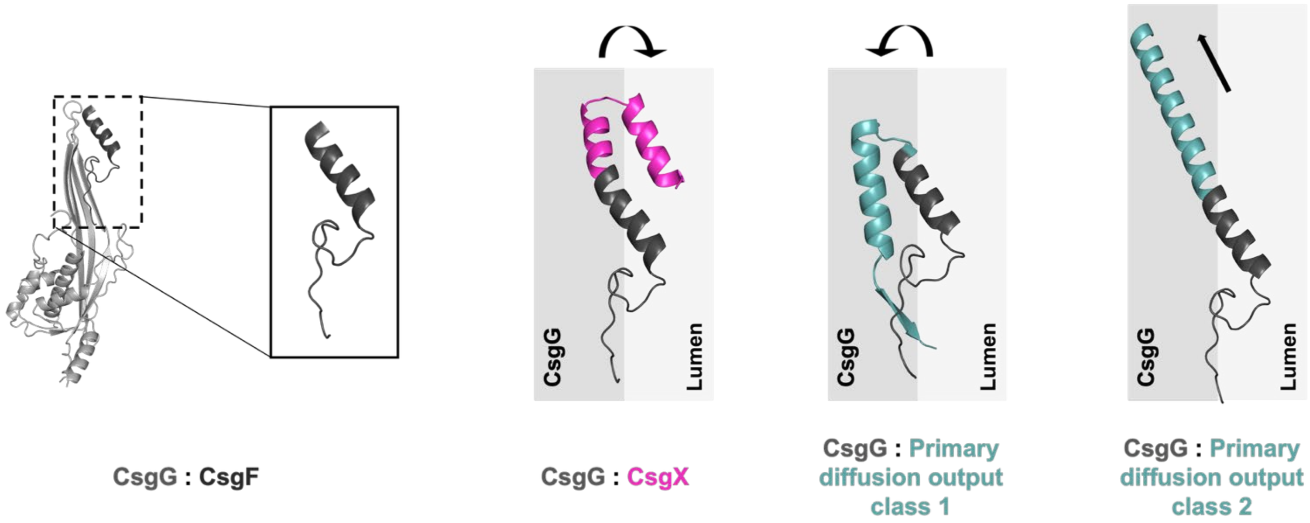


**Supplementary Data Figure 12: Attempts to install CsgX extension using RFdiffusion.** Multiple unsuccessful attempts to scaffold CsgX-like topologies were made. Two main classes of output were identified: either elongations folded towards CsgG and not the lumen (class 1), or further elongation of the Anchor (class 2). Although this might be possible through more careful tuning of parameters, we instead sought an approach that would allow us to design and evaluate interfaces that are as close to optimal as possible given the data available in the PDB, while adding transparency to enable the extraction of physical principals for construction of functional nanopores.

### Supplementary Data Table 1: CsgG, CsgF and CsgX sequences

| Design ID | Primary sequence | Outcome |
|---|---|---|
| CsgG-F56Q-StrepII | CLTAPPKEAARPTLMPRAQSYKDLTHLPAPTGKIFVSVYNIQDETGQFKPYPASNQSTAVPQSATAMLVTALKDSRWFIPLERQGLQNLLNERKIIRAAQENGTVAINNRIPLQSLTAANIMVEGSIIGYESNVKSGGVGARYFGIGADTQYQLDQIAVNLRVVNVSTGEILSSVNTSKTILSYEVQAGVFRFIDYQRLLEGEVGYTSNEPVMLCLMSAIETGVIFLINDGIDRGLWDLQNKAERQNDILVKYRHMSVPPESSAWSHPQFEK | Used for all (CsgG:CsgX)$_9$ assemblies in experimental and structural characterization |
| CsgF 1-30 | GTMTFQFRNPNFGGNPNNGAFLLNSAQAQN | Pore assembly, gel and electrophysiological analysis |
| CsgX1 | GTMTFQFRNPNFGGNPNNGAFLLNSAQAQNAGILAAQLWNNGDYDRALSLFIAVVQS-NH$_2$ | Full experimental and structural characterization |
| CsgX1$_{cyc}$ | GTMTFQFRNPNFGGNPNNGAFLL**C**SAQAQNAGILAAQLWNNGDYDRALSLFIAVVQS**C**-NH$_2$ | Full experimental and structural characterization |
| CsgX2 | GTMTFQFRNPNFGGNPNNGAFLLNSAQAQNGGELAAKLWANGDETNALSLFQTIIQS-NH$_2$ | Full experimental and structural characterization |
| CsgX2$_{cyc}$ | GTMTFQFRNPNFGGNPNNGAFLL**C**SAQAQNGGELAAKLWANGDETNALSLFQTIIQS**C**-NH$_2$ | Full experimental and structural characterization |
| CsgX3 | GTMTFQFRNPNFGGNPNNGAFLLNSAQAQNAGELAKKLWENGNVNQALSLFQTVIQS-NH$_2$ | Experimental characterization |
| CsgX3$_{cyc}$ | GTMTFQFRNPNFGGNPNNGAFLL**C**SAQAQNAGELAKKLWENGNVNQALSLFQTVIQS**C**-NH$_2$ | Experimental characterization |
| CsgX4 | GTMTFQFRNPNFGGNPNNGAFLLNSAQAQNAGELGLKLLRKGDVETALTLFAQVISG-NH$_2$ | Pore assembly |
| CsgX4$_{cyc}$ | GTMTFQFRNPNFGGNPNNGAFLL**C**SAQAQNAGELGLKLLRKGDVETALTLFAQVISG**C**-NH$_2$ | Pore assembly |
| CsgX5 | GTMTFQFRNPNFGGNPNNGAFLLNSAQAQNAGELGLKLLRKGDVETALKLFAIVIAG-NH$_2$ | Pore assembly |
| CsgX5$_{cyc}$ | GTMTFQFRNPNFGGNPNNGAFLL**C**SAQAQNAGELGLKLLRKGDVETALKLFAIVIAG**C**-NH$_2$ | Pore assembly |

**Supplementary Data Table 2: Cryo-EM validation data**

| | $(CsgG:CsgX1)_9$ | $(CsgG:CsgX1_{cyc})_9$ | $(CsgG:CsgX2)_9$ | $(CsgG:CsgX2_{cyc})_9$ |
|---|---|---|---|---|
| **Data collection and processing** | | | | |
| Microscope and camera | Titan Krios, K3 | | | |
| Magnification | 105,000 | | | |
| Voltage | 300 | | | |
| Data acquisition software | Serial EM | | | |
| Exposure navigation | Image shift | | | |
| Electron exposure ($e^-$/Å$^2$) | 46 | | | |
| Exposure per frame (sec) | 0.024 | | | |
| Defocus range (µm) | -0.8 to -1.8 | | | |
| Movies collected | 2,886 | 2,299 | 2,887 | 2,745 |
| Box size | 440 | | | |
| Symmetry Imposed | C9 | C1 | C9 | C9 |
| Final particle images (no.) | 79,368 | 784,377 (symmetry expanded from 97,153 with C9 symmetry) | 77,499 | 100,419 |
| Map resolution (Å) | 2.59 | 2.84 | 2.62 | 2.57 |
| B factor (Å$^2$) | -103.5 | -93.8 | -99.7 | -104.1 |
| FSC threshold | 0.143 | | | |
| Map resolution range (Å) | 2.2–3.1 | 2–20 | 2.2–3.6 | 2.2–3.1 |
| **Refinement** | | | | |
| Model resolution (Å) | 2.6 | 2.7 | 2.6 | 2.5 |
| FSC threshold | 0.143 | | | |
| **Model composition** | | | | |
| Non-hydrogen atoms | 21,618 | 21,654 | 19,260 | 21,096 |
| Protein residues | 2790 | 2799 | 2484 | 2736 |
| Ligands | n/a | n/a | n/a | n/a |
| **B factors (Å$^2$)** | | | | |
| Protein | 86.27 | 121.21 | 80.35 | 77.78 |
| Ligand | n/a | n/a | n/a | n/a |
| **R.m.s. deviations** | | | | |
| Bond lengths (Å) | 0.002 | 0.004 | 0.003 | 0.004 |
| Bond angles (°) | 0.500 | 0.513 | 0.575 | 0.561 |
| **Validation** | | | | |
| Molprobity score | 0.72 | 0.98 | 1.04 | 0.85 |
| Clashscore | 0.67 | 2.11 | 2.55 | 1.28 |
| Rotamer outliers (%) | 0.39 | 0.26 | 0.43 | 0.00 |

### Supplementary scripts:

### MASTER search commands:

**Backbone Search:**
~/Path/master --targetList ~/Path/pds/list --rmsdCut 1 --query query.pds --matchOut ./output/match.txt --structOut ./output/ --outType full --seqOut ./output/seq.txt --topN 200
**Master Designability Analysis:**
~/Path/master --targetList ~/Path/pds/list --rmsdCut 1.5 --query query.pds --matchOut ./output/match.txt --structOut ./output/ --outType match --seqOut ./output/seq.txt

**Master Looping:**
~/Path/master --targetList ~/Path/pds/list --query query.pds --rmsdCut 1 --gapLen 8 21 --matchOut ./output/match.txt --structOut ./output/ --outType wgap --topN 200

### Example python script for Master Looping and Clustering:

```
## code to iterate the MASTER loop searching and clustering automatically. This file needs to be in the directory containing the .pdb of interest.
# Example syntax:
#"python MASTER_loop_search.py 8 21 1" will search for gapLen's starting from 8 and will run the wgap until length 21 with RMSD 1 and then perform loop clustering.
# Change the --targetList as needed

import numpy as np
import sys
import math
import os
import prody as pr
from scipy.sparse import csr_matrix

# Check if an "output" directory exists, make if not present.
dir_path = os.path.dirname(os.path.realpath(__file__))
isdir = os.path.isdir(dir_path + '/output')
print(isdir)
if isdir == False:
    print("'output' directory does not exist; making new directory...")
    directory = "output"
    os.mkdir(directory)
    print("Directory '%s' created" %directory)

query=(sys.argv[1])

# Use the 'createPDS' module of MASTER on the query.pdb and grab the query.pds. Change the directory path as needed for your installation.
os.system("~/Path/createPDS --type query --pdb %s" % query)
```

```
name, extension=os.path.splitext(query)
pds= name + '.pds'

# Use the 'master' module of MASTER on the query.pds. Change the directory paths as needed for
your installation (i.e. targetList and MASTER modules)
start=int(sys.argv[2]) # grabs the initial starting gapLen
extend=int(sys.argv[3]) # grabs the length to query
rmsdCut=(sys.argv[4]) # grabs the rmsdCut per normal MASTER search
gapLen_list=list(range(start,(extend+1)))
for i in gapLen_list:
        gapLen=i
        os.system("~/Path/master --query %s --targetList ~/Path/pds/list --rmsdCut %s --gapLen %d --
matchOut ./output/%d/match.txt --structOut ./output/%d --outType wgap --topN 200" % (pds, rmsdCut,
gapLen, gapLen, gapLen))

## Runs the "cluster_loops.py" command:

def get_rot_trans(mob_coords, targ_coords):
    mob_coords_com = mob_coords.mean(0)
    targ_coords_com = targ_coords.mean(0)
    mob_coords_cen = mob_coords - mob_coords_com
    targ_coords_cen = targ_coords - targ_coords_com
    cov_matrix = np.dot(mob_coords_cen.T, targ_coords_cen)
    U, S, Wt = np.linalg.svd(cov_matrix)
    R = np.dot(U, Wt)
    if np.linalg.det(R) < 0.:
        Wt[-1] *= -1
        R = np.dot(U, Wt)
    return R, mob_coords_com, targ_coords_com


#@jit("f4[:,:](f4[:,:,:])", nopython=True, cache=True)
def _make_pairwise_rmsd_mat(X):
    M = X.shape[0]
    N = X.shape[1]
    O = X.shape[2]
    D = np.zeros((M, M), dtype=np.float32)
    m_com = np.zeros(O, dtype=np.float32)
    t_com = np.zeros(O, dtype=np.float32)
    m = np.zeros((N, O), dtype=np.float32)
    mtrans = np.zeros((O, N), dtype=np.float32)
    mtr = np.zeros((N, O), dtype=np.float32)
    t = np.zeros((N, O), dtype=np.float32)
    c = np.zeros((O, O), dtype=np.float32)
    U = np.zeros((O, O), dtype=np.float32)
    S = np.zeros(O, dtype=np.float32)
    Wt = np.zeros((O, O), dtype=np.float32)
    R = np.zeros((O, O), dtype=np.float32)
    mtr_re = np.zeros(N * O, dtype=np.float32)
    t_re = np.zeros(N * O, dtype=np.float32)
    sub = np.zeros(N * O, dtype=np.float32)
```

```
    for i in range(M):
        for j in range(i + 1, M):
            for k in range(O):
                m_com[k] = np.mean(X[i, :, k])
                t_com[k] = np.mean(X[j, :, k])
            m = np.subtract(X[i, :, :], m_com)
            for a in range(N):
                for b in range(O):
                    mtrans[b, a] = m[a, b]
            t = np.subtract(X[j, :, :], t_com)
            c = np.dot(mtrans, t)
            U, S, Wt = np.linalg.svd(c)
            R = np.dot(U, Wt)
            if np.linalg.det(R) < 0.0:
                Wt[-1, :] *= -1.0
                R = np.dot(U, Wt)
            mtr = np.add(np.dot(m, R), t_com)
            q = 0
            for a in range(N):
                for b in range(O):
                    mtr_re[q] = mtr[a, b]
                    t_re[q] = X[j, :, :][a, b]
                    q += 1
            sub = np.subtract(mtr_re, t_re)
            D[i, j] = np.sqrt(1.0 / N * np.dot(sub, sub))
    return D


class Cluster:
    def __init__(self, **kwargs):

        self.rmsd_cutoff = kwargs.get('rmsd_cutoff', 0.5)
        self.cluster_tag = kwargs.get('cluster_tag', '')
        self.selection = kwargs.get('selection', 'all')
        od = kwargs.get('rmsd_mat_outdir', '.')
        self.rmsd_mat_outdir = od if od[-1] == '/' else od + '/'
        od = kwargs.get('clusters_outdir', '.')
        self.clusters_outdir = od if od[-1] == '/' else od + '/'
        self.pdbs = None
        self.pdb_coords = list()
        self.pdbs_errorfree = list()
        self.rmsd_mat = None
        self.adj_mat = None
        self.mems = None
        self.cents = None
        self._square = False
        self._adj_mat = False
        self._data_set = False
        self._ifg_count = list()
        self._vdm_count = list()
        self._query_name = list()
```

```
        self._centroid = list()
        self._cluster_num = list()
        self._cluster_size = list()
        self._rmsd_from_centroid = list()
        self._cluster_type = list()
        self.df = None

    def set_pdbs(self, pdbs):
        """Load pdbs into the Cluster object to be clustered.
        VdMs must be all one residue type."""
        self.pdbs = pdbs

    def _set_coords(self, pdb):
        """Grabs coordinates of atoms in bb selection and iFG dict."""
        try:
            coords = pdb.select(self.selection).getCoords()
            self.pdb_coords.append(coords)
            self.pdbs_errorfree.append(pdb)
        except AttributeError:
            pass

    def set_coords(self):
        """Grabs coords for every vdM in pdbs."""
        for pdb in self.pdbs:
            self._set_coords(pdb)
        self.pdb_coords = np.array(self.pdb_coords, dtype='float32')

    def make_pairwise_rmsd_mat(self):
        """Uses C-compiled numba code for fast pairwise superposition
        and RMSD calculation of coords (all against all)."""
        assert isinstance(self.pdb_coords, np.ndarray), 'PDB coords must be ' \
                                                        'numpy array'
        assert self.pdb_coords.dtype == 'float32', 'PDB coords must ' \
                                                   'be dtype of float32'
        self.rmsd_mat = _make_pairwise_rmsd_mat(self.pdb_coords)

    @staticmethod
    def greedy(adj_mat):
        """Takes an adjacency matrix as input.
            All values of adj_mat are 1 or 0:  1 if <= to cutoff, 0 if > cutoff.
            Can generate adj_mat from data in column format with:
            sklearn.neighbors.NearestNeighbors(metric='euclidean',
            radius=cutoff).fit(data).radius_neighbors_graph(data)"""

        if not isinstance(adj_mat, csr_matrix):
            try:
                adj_mat = csr_matrix(adj_mat)
            except:
                print('adj_mat distance matrix must be scipy csr_matrix '
                      '(or able to convert to one)')
                return
```

```
        assert adj_mat.shape[0] == adj_mat.shape[1], 'Distance matrix is not square.'

        all_mems = []
        cents = []
        indices = np.arange(adj_mat.shape[0])

        while adj_mat.shape[0] > 0:
            cent = adj_mat.sum(axis=1).argmax()
            cents.append(indices[cent])
            row = adj_mat.getrow(cent)
            tf = ~row.toarray().astype(bool)[0]
            mems = indices[~tf]
            all_mems.append(mems)
            indices = indices[tf]
            adj_mat = adj_mat[tf][:, tf]

        return all_mems, cents

    def make_square(self):
        self.rmsd_mat = self.rmsd_mat.T + self.rmsd_mat
        self._square = True

    def make_adj_mat(self):
        """Makes an adjacency matrix from the RMSD matrix"""
        self.adj_mat = np.zeros(self.rmsd_mat.shape)
        self.adj_mat[self.rmsd_mat <= self.rmsd_cutoff] = 1
        self.adj_mat = csr_matrix(self.adj_mat)
        self._adj_mat = True

    def cluster(self):
        """Performs greedy clustering of the RMSD matrix with a given
        RMSD cutoff (Default cutoff is 0.5 A)."""
        assert self.rmsd_mat is not None, 'Must create rmsd matrix first with ' \
                                          'make_pairwise_rmsd_mat()'
        if not self._square:
            self.make_square()

        if not self._adj_mat:
            self.make_adj_mat()

        self.mems, self.cents = self.greedy(self.adj_mat)

    def print_cluster(self, cluster_number, outpath=None):
        """Prints PDBs of a cluster after superposition of the backbone (bb_sel)
        onto that of the cluster centroid.  The backbone of the cluster centroid
        is itself superposed onto that of the largest cluster's centroid."""

        if not outpath:
            outpath = self.clusters_outdir + '/' + str(cluster_number) + '/'
```

```
        try:
            os.makedirs(outpath)
        except FileExistsError:
            pass

        cluster_index = cluster_number - 1
        cent = self.cents[cluster_index]
        mems = self.mems[cluster_index]

        # Align backbone of cluster centroid to backbone of centroid of largest cluster.
        R, m_com, t_com = get_rot_trans(self.pdb_coords[cent],
                                        self.pdb_coords[self.cents[0]])
        cent_coords = np.dot((self.pdb_coords[cent] - m_com), R) + t_com

        for i, mem in enumerate(mems):
            R, m_com, t_com = get_rot_trans(self.pdb_coords[mem], cent_coords)
            pdb = self.pdbs_errorfree[mem].copy()
            pdb_coords = pdb.getCoords()
            coords_transformed = np.dot((pdb_coords - m_com), R) + t_com
            pdb.setCoords(coords_transformed)
            is_cent = '_centroid' if mem == cent else ''
            pr.writePDB(outpath + 'cluster_' + str(cluster_number) + '_mem_' + str(i)
                        + self.cluster_tag + is_cent + '_' + str(pdb).split()[-1] + '.pdb.gz', pdb)

    def print_clusters(self, clusters):
        """Prints all clusters in the list *clusters*"""
        for cluster_number in clusters:
            self.print_cluster(cluster_number)

    def run_protocol(self, pdbs):
        """Runs a series of methods that will cluster the
        PDBs and output a pandas dataframe, RMSD matrix, and
        PDBs of top 20 clusters."""
        print('begin clustering...')
        self.set_pdbs(pdbs)
        print('setting PDB coords...')
        self.set_coords()
        print('constructing RMSD matrix...')
        self.make_pairwise_rmsd_mat()
        print('greedy clustering...')
        self.cluster()
        print('making dataframe...')
        self.print_clusters(clusters=range(1, len(self.mems) + 1))


def listdir_mac(path):
    return [f for f in os.listdir(path) if f[0] != '.']


##########
##########
```

```
def run_cluster(path):
    for loopdir in listdir_mac(path):
        print(loopdir)
        loopsize = int(loopdir)
        pdbs = list()
        for f in [f for f in os.listdir(path + loopdir) if f[-3:] == 'pdb']:
            pdb = pr.parsePDB(path + loopdir + '/' + f)
            ca_sel = pdb.select('name CA')
            if len(ca_sel) == 14 + loopsize:
                pdbs.append(pdb)

        kwargs = dict(rmsd_cutoff=1.0, cluster_tag='_loop_'+str(loopsize),
                      clusters_outdir=path + loopdir + '/clusters/',
                      selection='name CA')
        clu = Cluster(**kwargs)
        clu.run_protocol(pdbs)

path = dir_path + '/output/'
run_cluster(path)
print("Loop outputs and clusters can be found in '%s'" %path)
```

**Rosetta sequence design file:**

```
<ROSETTASCRIPTS>
                    <SCOREFXNS>
                         <ScoreFunction name="ref15" weights="ref2015">
                         </ScoreFunction>
                    </SCOREFXNS>

                    <RESIDUE_SELECTORS>
                    </RESIDUE_SELECTORS>

                    <TASKOPERATIONS>
                         <InitializeFromCommandline name="ifcl"/>
                         <ReadResfile name="resfile" filename="resfile.txt"/>
                         <ExtraRotamersGeneric name="extrachi" ex1="1" ex2="1"
ex1_sample_level="1" ex2_sample_level="1" extrachi_cutoff="14"/>
                         <IncludeCurrent name="include_curr" />
                    </TASKOPERATIONS>

                    <FILTERS>
                         <PackStat name="pstat" threshold="0.30" repeats="10" />
                         <PackStat name="pstat_mc" threshold="0" repeats="10"/>
                         <NetCharge name="net_charge" confidence="0"/>
                         <ScoreType name="total_score" scorefxn="ref15"
score_type="total_score" threshold="100000000000"/>
                         <ScoreType name="total_score_1" scorefxn="ref15"
score_type="total_score" threshold="100000000000"/>
                    </FILTERS>

                    <MOVERS>
                         <SetupForSymmetry name="setup_symm"
definition="spONT.symm"/>

                         <PackRotamersMover name="pack" scorefxn="ref15"

task_operations="ifcl,resfile,include_curr,extrachi"/>
                         <PackRotamersMover name="pack_fast" scorefxn="ref15"

task_operations="ifcl,resfile,include_curr"/>
                         <MinMover name="min_bb" scorefxn="ref15"
tolerance="0.0000001" max_iter="1000" chi="false" bb="true">
                              <MoveMap name="map_bb">
                                   <Span begin="1" end="999" bb="true" chi="false"
/>

                                   Span begin="120" end="999" bb="true"
chi="false"/>

                                   Jump number="2" setting="1" />
                              </MoveMap>
                         </MinMover>
                         <Idealize name="idealize"/>
                         <MinMover name="min_sc" scorefxn="ref15"
tolerance="0.0000001" max_iter="1000" chi="true" bb="false">
```

```
				<MoveMap name="map_sc">
					<Span begin="1" end="999" bb="false" chi="true"
/>

				</MoveMap>
			</MinMover>
			<MinMover name="min_sc_bb" scorefxn="ref15"
tolerance="0.0000001" max_iter="1000" chi="true" bb="true">
				<MoveMap name="map_sc_bb">
					<Span begin="1" end="999" bb="true" chi="true" />

				</MoveMap>
			</MinMover>
			<ParsedProtocol name="parsed_pack_fast" >
				<Add mover_name="pack_fast"/>
				<Add mover_name="min_bb"/>
			</ParsedProtocol>
			<ParsedProtocol name="parsed_pack" >
				<Add mover_name="pack"/>
				<Add mover_name="min_bb"/>
				<Add mover_name="min_sc"/>
			</ParsedProtocol>
			<GenericMonteCarlo name="pack_mc" preapply="0" trials="3"
temperature="0.03"

	filter_name="pstat_mc" sample_type="high" mover_name="parsed_pack">
					<Filters>
						<AND filter_name="total_score_1"
temperature="15" sample_type="low"/>
					</Filters>
			</GenericMonteCarlo>
			<GenericMonteCarlo name="pack_fast_mc" preapply="0"
trials="2" temperature="0.03"

	filter_name="pstat_mc" sample_type="high" mover_name="parsed_pack_fast">
					<Filters>
						<AND filter_name="total_score_1"
temperature="15" sample_type="low"/>
					</Filters>
			</GenericMonteCarlo>

		</MOVERS>
		<APPLY_TO_POSE>
		</APPLY_TO_POSE>

		<PROTOCOLS>
			<Add mover="setup_symm"/>
			<Add mover_name="parsed_pack_fast"/>
			<Add mover_name="pack_fast_mc"/>
			<Add mover_name="pack_mc"/>
			<Add mover_name="min_sc_bb"/>
```

```
                <Add filter_name="pstat"/>
                <Add filter_name="net_charge"/>
                <Add filter_name="total_score"/>
        </PROTOCOLS>

        <OUTPUT scorefxn="ref15"/>
</ROSETTASCRIPTS>
```

**Base Rosetta Resfile:**

```
NATRO
USE_INPUT_SC
start

31 A PIKAA AG
32 A PIKAA G
34 A PIKAA LI
35 A PIKAA AG
38 A PIKAA L
47 A PIKAA A
48 A PIKAA L
50 A PIKAA L
51 A PIKAA F
54 A PIKAA VIA
55 A PIKAA VI
```
